# Analysis and design of disordered polypeptides with optimized sequence patterning properties

**DOI:** 10.64898/2026.02.20.707115

**Authors:** Arjun Singh, Ali I Ukperaj, Gabriel F Porto, Gregory L Dignon

**Affiliations:** Department of Chemical and Biochemical Engineering, Rutgers University, Piscataway, NJ, United States

## Abstract

Intrinsically disordered proteins (IDPs) exhibit phase separation behavior that is closely linked to their degree of single-chain compaction, which in turn is governed by both amino acid composition and sequence patterning. Existing metrics such as sequence charge decoration (SCD) and sequence hydropathy decoration (SHD) describe these effects but are largely limited to describing differences between sequences of similar length and overall composition. In this work, we present a shuffle-based normalization scheme for SCD and SHD, enabling comparison of sequence patterning between very different IDP sequences. Leveraging this normalization scheme toward design space, we develop a Monte Carlo based sequence design algorithm that generates novel IDPs with desired patterning features. Our design framework is further strengthened by incorporating additional metrics such as sequence aromatic decoration (SAD), compositional RMSD, and a previously developed sequence based ΔG predictor. We validate our approach through coarse-grained MD simulations, showing that the designed sequences exhibit tunable phase behavior. This strategy lays the groundwork for rational design of IDPs for biomedical and biotechnology applications, as well as basic biophysical research.

**Author summary:** Intrinsically disordered proteins behave similar to polymers in solution, having no defined structure. Their behavior is dictated by the collection of shapes the protein adopts, known as it’s “conformational ensemble” which is tuned by its amino acid sequence and the solution environment. In this work, we have developed parameters to describe the patterning of charged and hydrophobic amino acids within these protein sequences, which are predictive of their ability to phase separate and form dense liquid-like droplets in solution. Importantly, the parameters we develop are motivated by physics and can be applied across a large number of amino acid sequences rapidly. This will enable researchers to rapidly predict the behavior of large libraries of protein sequences. We have additionally developed a software to design randomized amino acid sequences with desired amino acid composition and patterning properties. Finally, we have tested our design scheme and parameters by running simulations of designed IDP sequences and quantified each of their ability to phase separate.

## Introduction

Biomolecular condensates are specialized cellular compartments that form without membranes, helping cells organize important molecules through a process called liquid-liquid phase separation (LLPS) [1]. Unlike traditional organelles, condensates form membraneless organelles, compartments which can assemble and disassemble in response to changing conditions and protein properties [2]. They play essential roles in organizing and regulating biological processes, such as DNA replication and repair, transcription, chromatin remodeling, RNA metabolism, ribosome biogenesis, protein quality control, cell division, intercellular adhesion, and signal transduction [3–9]. Disruptions in condensate dynamics have been linked to neurodegenerative diseases, cancer, and viral infections, with aberrant signaling and assembly potentially contributing to pathological states and pathways such as uncontrolled proliferation and aggregation [10–13]. Given their broad functional importance, understanding how biomolecular condensates form and operate is vital for unraveling their roles in both normal physiology and disease [14].

A major factor influencing condensate formation is the distinct behavior of intrinsically disordered proteins (IDPs), which lack a stable three-dimensional structure and instead fluctuate among a diverse ensemble of conformations [15]. This conformational ensemble is governed by the IDP’s amino acid sequence and composition, which in turn determines how it interacts with other molecules and contributes to phase separation and condensate dynamics [16, 17]. Compositional features such as net charge, mean hydropathy, and charge density offer valuable insights into the conformational preferences and phase behavior of IDPs [16, 18]. Beyond composition, the arrangement of charged or aromatic residues within the amino acid sequence further modulates chain compaction and phase separation propensity [19–22]. Understanding these principles opens the door to rational sequence design, not only to map how sequence governs behavior, but also to engineer IDPs for specific functions or subcellular localizations [23–26]. Moreover, short synthetic sequences can serve as minimal models for LLPS, enabling mechanistic studies under controlled conditions [27].

Computational and theoretical studies have shown that the degree of single-chain compaction in soluble IDPs strongly correlates with their phase separation propensity—more compact chains generally phase separate more readily [21, 28–30]. This compaction is influenced by overall protein composition as well as the patterning of particular amino acid chemistries within the sequence. For example, blocky distributions of like-charged or hydrophobic residues promote intra-chain attractions, leading to collapse, and in turn, enhance intermolecular interactions that drive LLPS [20, 21, 31]. Quantitative metrics such as sequence hydropathy decoration (SHD) and sequence charge decoration (SCD) capture these effects and help predict phase behavior [24, 32, 33]. Our goal here is to refine the definitions of various patterning parameters and utilize them in design of unique IDP sequences.

In this work, we address limitations in how sequence-based patterning parameters—such as sequence charge decoration (SCD) and sequence hydropathy decoration (SHD), are currently defined and applied. Previous studies have shown advances where modifications to a sequence charge patterning or arrangement of hydrophobic/aromatic residues has a significant impact on their phase behavior, validated by both simulations and experiments [34–36]. While these metrics capture how charged and hydrophobic residues are arranged along a sequence, it is difficult to compare the degree of “blockiness” between sequences that are of different length or composition. Thus, studies applying SCD and SHD are typically limited to mutagenesis or randomization of individual sequences. In this work, we look to address this issue by proposing a normalization scheme, akin to the z-score approach from Pappu lab’s NARDINI parameter set [37, 38] and further propose empirical methods of approximating the sampled distribution as a function of compositional terms. This method is then leveraged to develop a Monte Carlo algorithm for the directed design of novel sequences with desired patterning characteristics. To enhance the robustness of this design framework, we later incorporate additional parameters including sequence aromatic decoration (SAD), compositional RMSD, and a sequence-based ΔG predictor [39]. Finally, we validate the designed sequences across diverse protein sequences using HPS-Urry simulations, demonstrating the algorithm’s ability to generate proteins with tunable phase behavior, laying the groundwork for applications such as biomolecular condensate-based drug delivery and determination of compositional control within condensates [2, 40].

## Results

### Generalized parameters to quantify patterning within a fixed composition

Previous work has shown that the critical temperature of IDP phase separation can be roughly predicted by the degree of collapse as a single chain [21, 29, 30, 41], generally demonstrating that IDPs with a more collapsed conformation will be more capable of phase separating, while those more extended will be incapable of phase separation on their own. It has also been demonstrated that the patterning of charged and hydrophobic amino acids is predictive of IDP chain dimensions [19, 20, 32] and phase separation [21, 35, 36]. In this work, we focus specifically on the patterning parameters sequence hydropathy decoration (SHD) and sequence charge decoration (SCD). SHD is calculated from the scaled hydropathy values and sequence separation of all residues in an amino acid sequence:

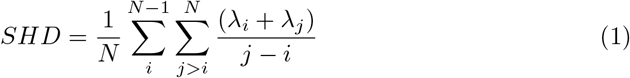

where N is the sequence length and *λ*_*i*_ is the scaled hydropathy value of residue i according to the Urry hydropathy scale [42] (S1 Table). SCD is calculated from the relative charge and sequence separation of formally charged residues in an amino acid sequence:

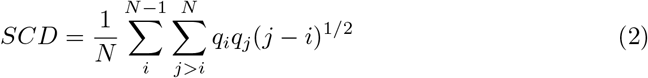

where N is the sequence length and q_*i*_ is the formal charge of residue i.

Both parameters are demonstrated to be very effective at predicting IDP conformations [32, 43] as well as phase separation and materials properties [33, 35]. We note, however, that there is a limitation to using such metrics since they effectively describe the degree of patterning, but only within sequences of identical length and composition. When used for sequences having drastically different composition or size, we observe widely varying values of SCD and SHD. To demonstrate this disconnect, we consider seven disordered sequences of different sizes and compositions from biology (S2 Table). To evaluate the effect of sequence composition on the range of accessible SCD and SHD values, we take an amino acid sequence and shuffle it 1 million times, producing only sequences with identical composition, but random ordering of amino acids. This can be considered as a representative ensemble of constant-composition sequences, from which we can approximate the underlying probability density of patterning parameters and the “blockiness” of sequences. From these sequence ensembles, we calculate SCD and SHD and plot the probability distributions (Fig. 1A,B). We note that for each individual sequence, the probability distributions have no overlap in SHD and only moderate overlap in SCD, making direct comparison of SHD or SCD values between different proteins challenging to interpret. For instance, the randomized sequence of FUS LC that is the least “blocky” has a greater SHD value than the most “blocky” randomized sequence of hnRNPA2 LC. Thus, we note the limitation of SCD and SHD for quantifying blockiness of disparate sequences, particularly those with differing size and composition, and highlight the need for a universal normalization scheme.

**Fig 1.**
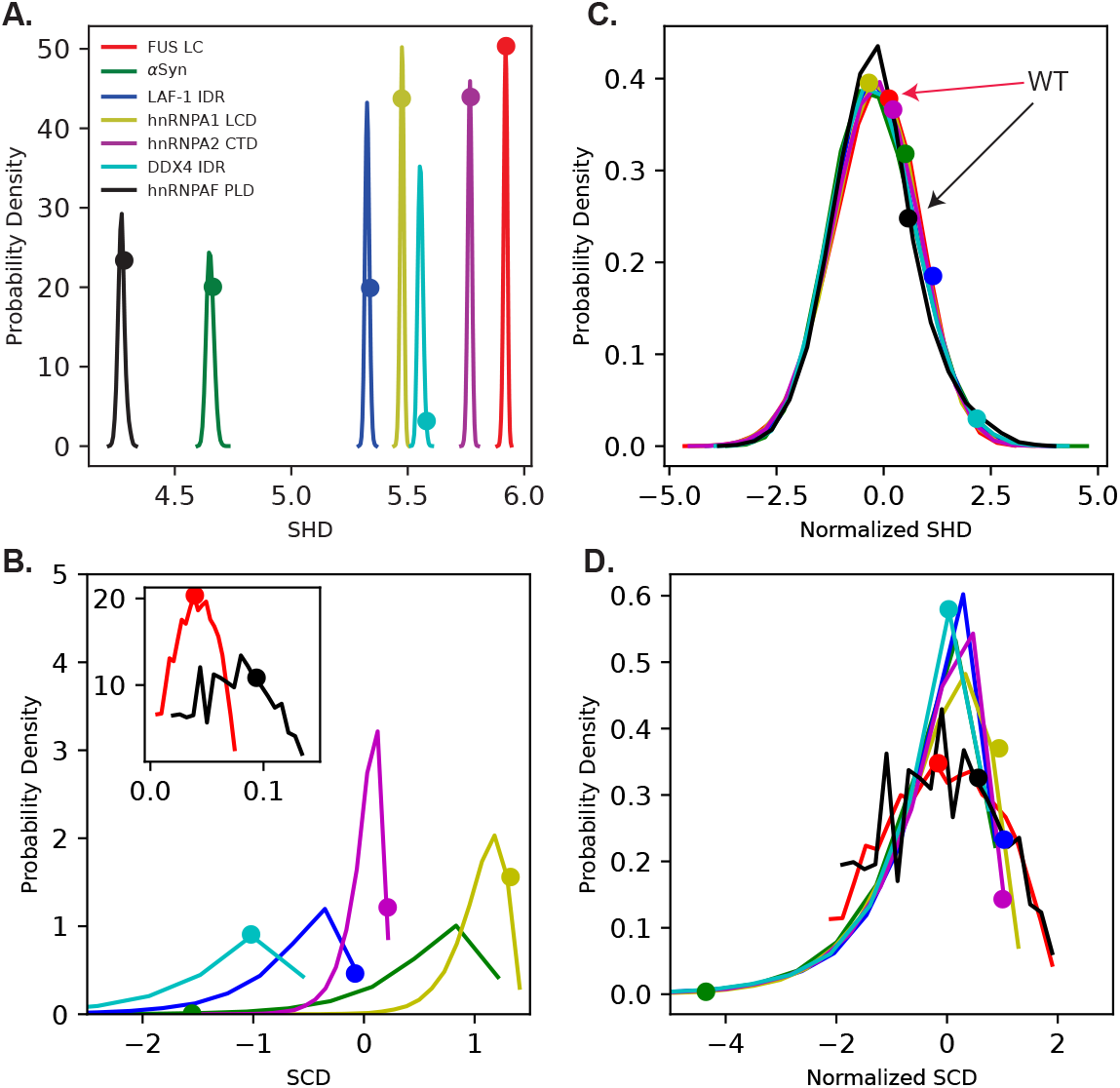
The probability density of patterning parameter values based on real IDP sequences that have been shuffled 1 million times each. A) SHD values of shuffled IDP sequences each follow normal distributions with different mean values, and relatively small spread. B) SCD values from shuffled sequences show more spread and overlap than SHD (FUS and hnRNPF shown in inset since they have very few charged residues), C) collapsed SHD_*norm*_ and D) SCD_*norm*_ values match well with the standard normal distribution, with SCD_*norm*_ having an extended tail toward negative numbers. In all panels, circles indicate the WT sequence value on each distribution.

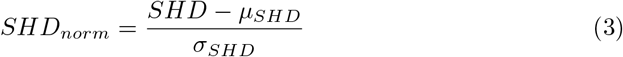

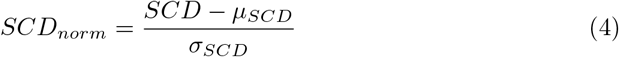

We find that the probability distributions are roughly gaussian in shape, allowing each distribution to be collapsed to the standard normal distribution (Fig. 1C,D). This normalization serves to identify the significance of a value within each parameter’s respective fixed-composition distributions, giving a sense of whether this protein sequence is well-mixed or segregated for something of its composition. For example, the *SCD*_*norm*_ of WT LAF-1 RGG is on the right arm of the probability distribution, while WT *α*-synuclein is far to the left of the main density of the distribution. This indicates that the charged residues of LAF-1 RGG and *α*-synuclein are more evenly distributed and more blocky than would be expected if randomly arranged, respectively. We note that while raw SCD values tend to have wider overlapping distributions and SHD has narrow, non-overlapping distributions, this is simply due to the number of sample distance comparisons within the given sequences, having all residue pairs included in SHD calculations, but only charged residues considered for SCD calculations.

We then ask the question of whether we can bypass the need for shuffling sequences to obtain the probability density of accessible SCD and SHD values. Toward this goal of approximating *µ*_*SHD*_ and *σ*_*SHD*_ for a given composition, we must first identify descriptors that don’t change upon shuffling, being only composition-dependent and not sequence-dependent. Starting with SHD, we find that the shuffled ensemble probability distribution can be predicted by just three composition-based parameters, i.e. the mean hydropathy (commonly denoted *< λ >*, but referred to here as *µ*_*λ*_), standard deviation of hydropathy (*σ*_*λ*_) of all residues in the sequence, and chain length (N). To test the predictability, we designed 48 mock sequences ranging in size from 50 to 200 residues, with *µ*_*λ*_ between 0.4 to 0.7, and with *σ*_*λ*_ values up to 0.5 depending on what was reasonably attainable for the given *µ*_*λ*_. For example, a polyglycine sequence will have *µ*_*λ*_ ~ 0.5 and *σ*_*λ*_ = 0, while a sequence comprising half glutamate and half tryptophan will have roughly the same *µ*_*λ*_, but will have a larger *σ*_*λ*_ value ~ 0.5. Taking these 48 sequences, we shuffled each one million times, and calculated the *µ*_*SHD*_ and *σ*_*SHD*_ values to provide real distributions which can then be predicted by the composition-based parameters (Fig. 2).

**Fig 2.**
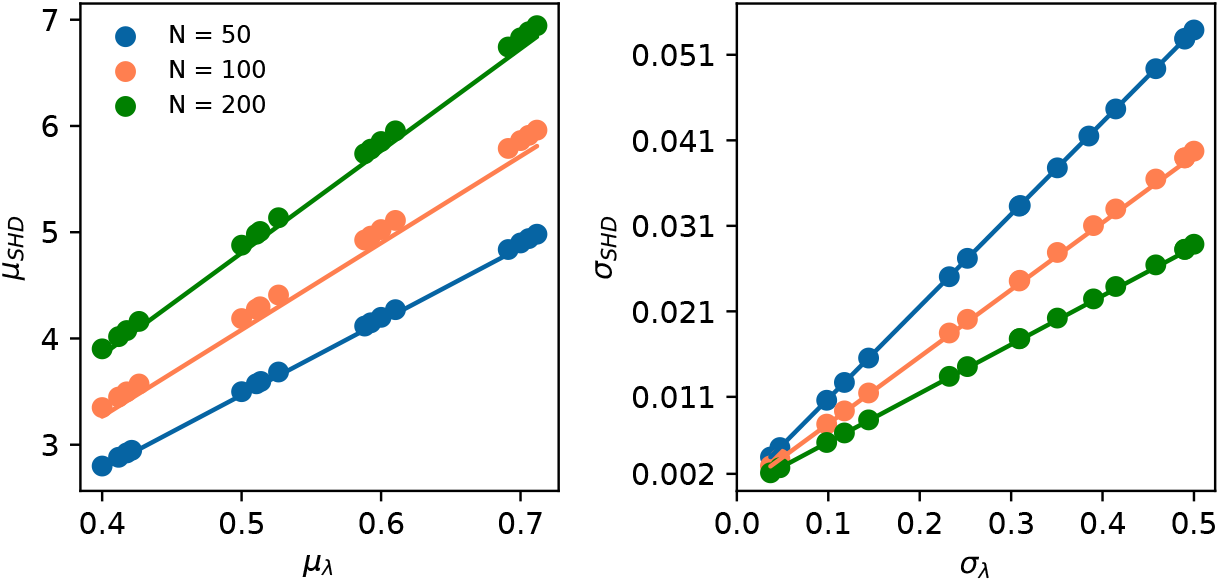
Predicting probability density of SHD from composition-dependent parameters. A) The average SHD for a constant-composition ensemble depends on the average hydropathy and length of the sequence. B) The standard deviation of SHD values for constant-composition ensemble depends on standard deviation of hydropathy and length.

We first compare *µ*_*SHD*_ with *µ*_*λ*_ and find that there is a positive linear correlation, having a slope dependent on the length of the sequence (Fig. 2A). We then compare *σ*_*SHD*_ with with *σ*_*λ*_ and find another linear correlation with an N-dependent slope (Fig. 2B). Notably, *µ*_*SHD*_ is larger for longer sequences, indicating that the value of SHD increases for longer and more hydrophobic sequences (Fig. 2A). In contrast, *σ*_*SHD*_ is smaller with increasing N, indicating smaller fluctuations occurring for greater sampling (Figure 2B). Intriguingly, we find that the average hydropathy is not predictive of *σ*_*SHD*_ on its own, and the standard deviation of hydropathy is not predictive of *µ*_*SHD*_ (S1 Fig). Using this data set, we derive a simple empirical formula for both the mean and standard deviation of possible SHD values based solely on sequence composition within the bounds of the tested sequences,

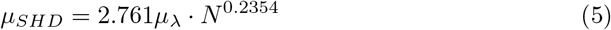

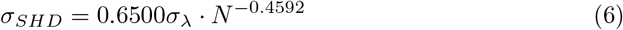

This enables prediction of the probability distribution of SHD values for any amino acid sequence without the need for hundreds of thousands of shuffles, and provides a useful normalization scheme for the SHD patterning parameter.

We repeated this process for the SCD parameter, aiming to identify composition-dependent but sequence-independent features that could be used to estimate *µ*_*SCD*_ and *σ*_*SCD*_. We similarly generated a library of synthetic sequences with varied charge compositions, having varied values of fraction of charged residues (FCR), net charge per residue (NCPR), and chain length (N). We observed that both the mean (*µ*_*SCD*_) and standard deviation (*σ*_*SCD*_) of the SCD distribution showed approximately linear relationships with FCR (Fig. 3A,B), indicating that FCR alone provides a useful predictor in these cases. On the other hand, the relationships between SCD statistics and NCPR were non-linear (Fig. 2C,D), suggesting more complex dependencies when considering net charge effects. Using these trends, we determined empirical equations that combine FCR, NCPR, and N to estimate the mean and standard deviation of the SCD distribution for a given sequence composition:

**Fig 3.**
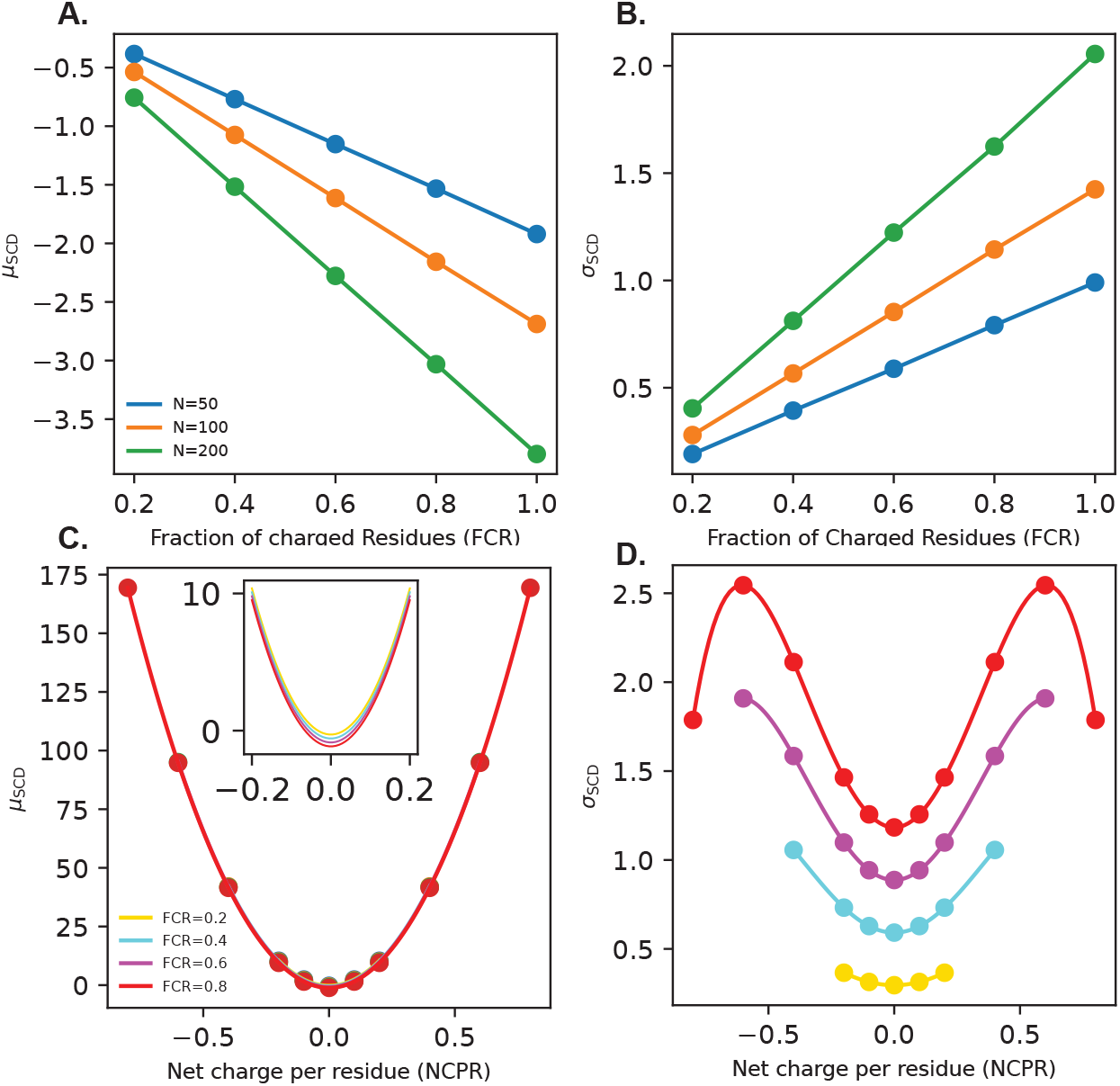
Prediction of SCD distribution from sequence-independent parameters. Each point represents statistics calculated from 1 million shuffled sequences per composition, with solid lines indicating empirical fits. A-B) Mean and standard deviation of SCD vs. FCR for sequences with net charge of zero. C-D) Mean and standard deviation of SCD vs. NCPR for sequences of size N=100. Inset in panel C highlights smaller influence of FCR on average SCD.

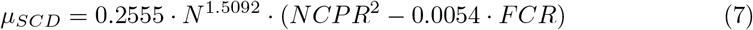

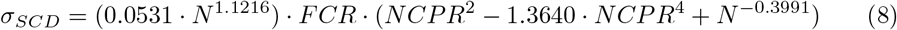

In addition to charge and hydropathy patterning, there is also known to be an influence of aromatic residue patterning on IDP conformations and phase separation [22]. We thus develop a third patterning parameter, focused specifically on aromatic residues within the amino acid sequence, termed sequence aromaticity decoration (SAD) and following a similar functional form to SHD:

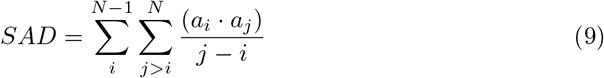

where *a*_*i*_ and *a*_*j*_ are the aromaticity of residues i and j. Aromatic residues are assigned a value of 1 and non-aromatic residues a value of 0. We find that upon the shuffling of random sequences, AD also follows a roughly gaussian distribution, and can be predicted using the following empirical equations:

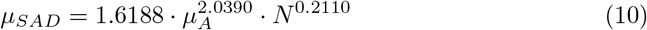

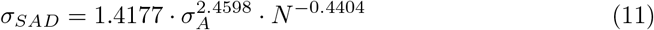

where *µ*_*SAD*_ and *σ*_*SAD*_ are the mean and standard deviation of aromaticity in the sequence, respectively. Intriguingly, both mean and standard deviation of the SAD distribution are dependent on both mean and standard deviation of the aromaticity in the amino acid sequence (*µ*_*A*_ and *σ*_*A*_). We find this is due to the fact that aromaticity is a binary in the sequence, and thus, the mean and standard deviation of aromaticity in a set composition are related to each other through the relation of random binary data, i.e. 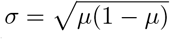, since *µ*_*A*_ is constrained between 0 and 1. SAD will serve as a useful design parameter as it should be able to help design sequences with well-dispersed aromatic residues, and reduce the likelihood of blocks of aromatic residues that can lead to aggregation or more solid or gel-like behavior of condensates [22]. We are also curious about the redundancy of these three parameters in describing IDP dimensions since each is dependent on a summation over sequence pair separation *j* − *i*, and somewhat correlated physicochemical properties of each amino acid (i.e. hydrophobic residues are not charged). To evaluate this, we generated 10^5^ random sequences with identical composition to 3 different IDPs used in this work, finding that the pairwise correlation of each pair of patterning parameters is quite minimal (S2 Fig). Indeed, the most significant Pearson correlation coefficient (r) is found between SHD and SAD in FUS LC with a value of 0.24.

### Sequence Monte Carlo to design polypeptides with desirable properties

Contextualization and normalization of patterning parameters across all compositions and sizes offers great potential to accelerate design of sequences with multiple optimized physical properties. We have effectively created a coordinate space within which values of these patterning parameters are equally weighted regardless of differing sequence compositions and lengths. Toward this goal, we leverage a Monte Carlo sampling algorithm to design new sequences with desirable properties. This includes quickly formulating sequences with targeted values of patterning parameters, as well as creating identical compositions of existing sequences with a tuned propensity to phase separate.

We first define a loss function as a Monte Carlo energy function by inputting “target” values for a set of parameters, namely the aforementioned patterning parameters, and then measure the deviation of a designed sequence from each target value. Notably, this will favor sequence modifications that move each parameter (i) closer to the target value (*i*_*target*_), and penalize modifications that move further away.

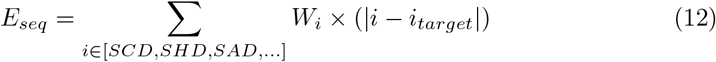

To ensure changes in each parameter are counted, and no one parameter is overly represented in the loss function, we set each parameter’s weight to *W*_*i*_ = 1*/σ*_*i*_, where *σ*_*i*_ is the standard deviation of that parameter’s probability distribution calculated with respect to the composition and using Equations 6, 8, and 11. This effectively makes the change in the MC energy equivalent to the sum of the change in *SCD*_*norm*_, *SHD*_*norm*_ and *SAD*_*norm*_. As a result of this weighting scheme, each parameter has roughly equivalent importance in the overall design and optimization of an IDP sequence. This approach can also incorporate other design parameters, such as aromatic patterning, and deviation from a set sequence composition (Fig. 4A-C). We can also incorporate direct predictions of phase separation capability, such as the Δ*G* predicted from sequence using the algorithm developed by von Büllow et al. [39]. For these terms, we are not able to produce similar weights as the previously described patterning terms, and thus opt to weigh them by their maximum possible value instead. We find that these weights keep term contributions within 2-3 orders of magnitude of changes to the patterning terms. We implement each of these and discuss further in the following sections. Notably, the process of sequence design is modular and can be made to incorporate even more design parameters than discussed in this work.

**Fig 4.**
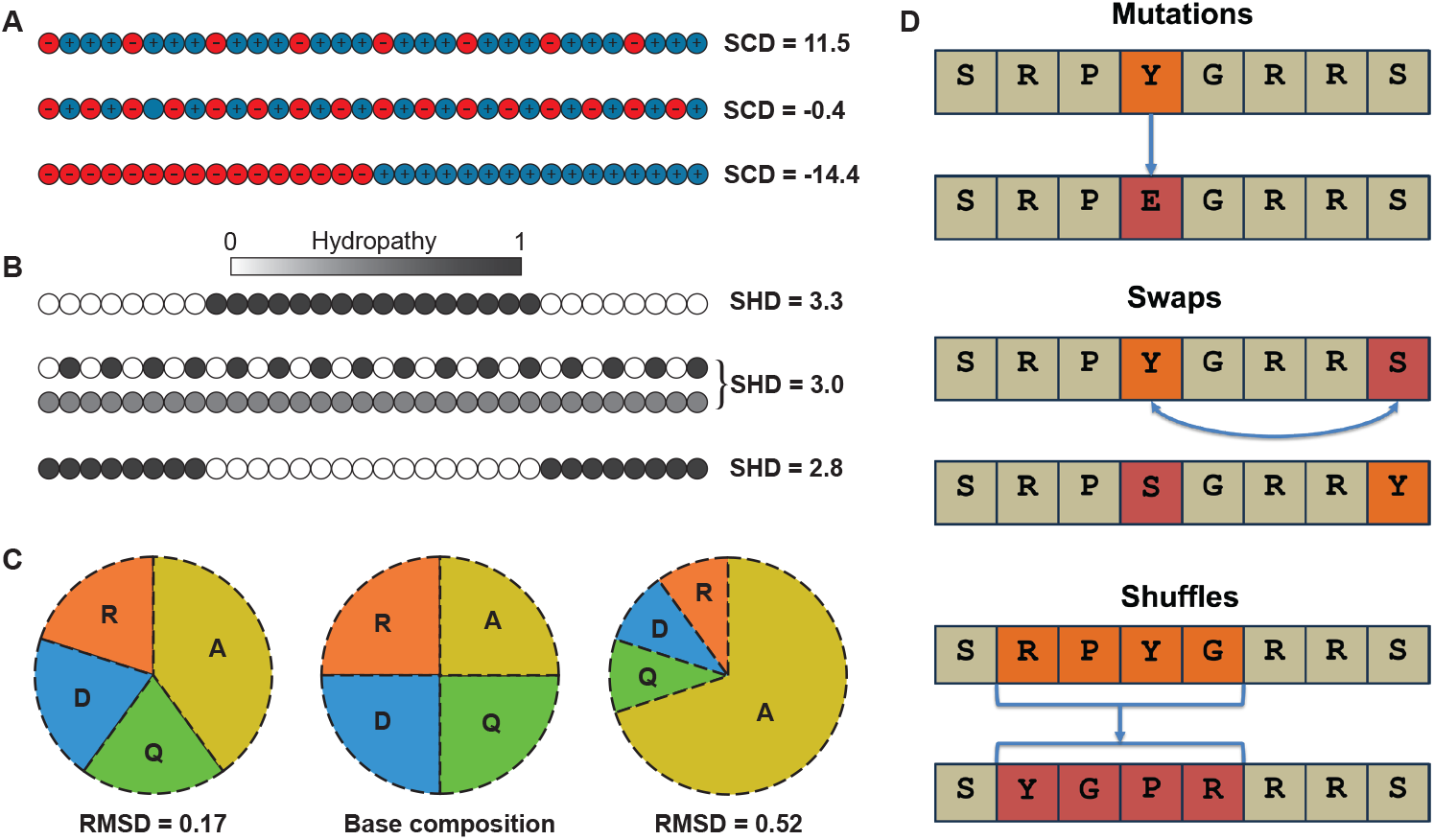
Design parameters and moveset for sequence Monte Carlo algorithm. A) SCD is related to the degree of segregation of charged residues, where well-mixed polyampholytes are close to zero, blocky polyampholytes have large negative SCD, and sequences with a significant net charge have a positive SCD. B) SHD tends to be higher for sequences having hydrophobic groups more localized toward the middle of the sequence. Typically, IDP sequences are not so extreme, so SHD functions to quantify blockiness of hydrophobic residues primarily, and is somewhat influenced by proximity of hydrophobic groups toward the edges of the sequence. C) Composition RMSD is a simple metric of how similar the current composition is from that of the base composition. D) Three types of moves are attempted for the MC code, mutation moves change a single residue’s type, swap moves swap the positions of two random residues, and shuffle moves randomize a part of the sequence.

To carry out sequence design, we use a Metropolis Monte Carlo approach where perturbations are made to the sequence, the new sequence “energy” is calculated, and moves are accepted or rejected based on the Metropolis Criterion [44]. To perturb the sequence, we introduce three types of “moves”, namely mutations, swaps and partial sequence shuffles (Fig. 4D). The MC moves result in changes in the loss function (Δ*E*) which is then used to accept or reject the move based on the Metropolis criterion

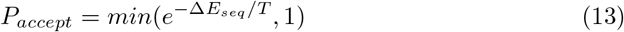

where T is a scaling parameter that determines the likelihood of accepting uphill moves to higher energies, potentially aiding the escape of local minima in the energy function. For our study, we have set this to be 10^−2^ as an intermediate value allowing some exploration, but ensuring overall migration toward the optimized design parameters. This process will repeat until all parameters are within an interval of their desired values, or until the maximum set number of desired cycles has been reached.

As a trial run, we use LAF-1 RGG as an input and begin to design a target sequence with each design parameter set to several standard deviations away from the initial value, with the exception of composition, which is set to stay as close to the original sequence composition as possible—RMSD = 0. SCD, SHD, and SAD optimization was particularly fast, reaching target values in around 200 moves (Fig. 5A,B). Alternatively, the ΔG predictor was able to reach its target value within an attempted 2000 Monte Carlo moves (Fig. 5C,D). The composition of the sequence was allowed to fluctuate, but only did so moderately (Fig. 5E), staying very close to the original sequence composition. Consequently, the total loss function reached a minimum at ~ 200 moves (Fig. 5F). It is important to note that, while these trajectories converged successfully to target values in a low number of moves, this success is only possible because targets were reasonably attainable given the input sequence. For targets well outside the range of the optimized parameter’s distribution for the given composition, the algorithm will be unsuccessful or take a significantly longer time to reach it through the gradual mutation of the sequence. It is also possible to optimize just a single parameter, such as SCD (Fig. S3), however we find that other parameters can widely vary and fluctuate throughout the design process (Fig. S9). Depending on one’s design goals, this may be disadvantageous.

**Fig 5.**
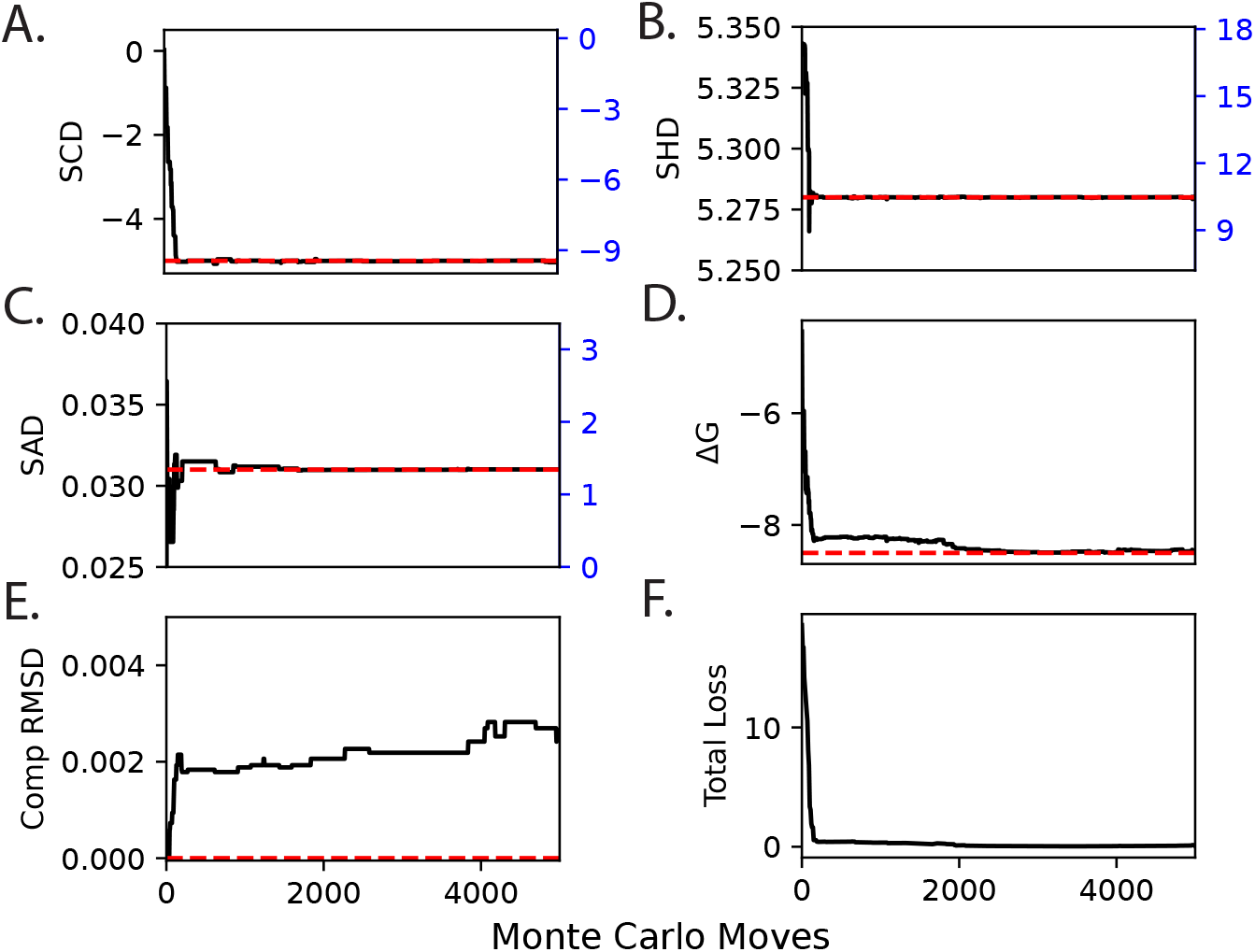
(A-E) Trajectory of sequence MC for LAF-1 RGG converging to desirable properties of SCD, SHD, SAD, ΔG, and Composition RMSD, respectively. Blue axes represent the z-score of the constant-composition distribution for the patterning parameters. (F) Total loss progression over 5000 Monte Carlo moves.

To evaluate the success of the normalization scheme for patterning parameters, we plotted the energy decomposition for additional design runs of LAF-1 RGG, breaking down the contributions of each parameter by what move type was attempted. In order to minimize drift throughout the course of the simulation, target values were set equal to the initial values of the input sequence (S2 Fig). Comparing between these parameters within a single move type, we can evaluate what each term’s average energy contribution was. In this manner, we would expect a successful normalization scheme to keep all contributions within a reasonable range of one another. We find that for SCD, SHD, and SAD, each move type only exhibits variations within an order of magnitude of energy fluctuations (S2 Fig), indicating successful normalization of these parameters.

### Efficient and guided design of extreme patterning in sequences

With the identification of significant changes in phase-behavior between protein sequences of fixed-composition, a commonly explored area of interest is the relation between patterning parameters and a protein’s propensity for phase separation. In a previous study, we investigated the relationship between the phase-behavior of fixed-composition LAF-1 RGG variants at a wide-range of SCD values [35]. By performing 10 million random shuffles to the wild type sequence with an SCD of 0.6, the variant with the lowest attained SCD only reached −7.3. To recreate this effort using our Monte Carlo approach, we utilize LAF-1 RGG as a reference sequence to which further variants will be designed through shuffling. To keep composition from changing, we disabled mutation moves for this set of sequence designs. The shuffling moves attempted differ from the original study in that they are applied to only segments of the sequence, and their contribution to SCD is evaluated, such that changes that increase SCD are not accepted. This causes the sequence to walk a directed path toward the desired properties. We repeat this process 10 times for LAF-1 RGG variants with three different target SCDs of −5, −10, and −15, which averaged 180, 370, and 800 cycles, respectively, to complete (Fig. 6A-C). Each design task, two of which extended significantly past the attainable sequences from the aforementioned paper, was attained in fewer than 1000 shuffle moves. Notably, the calculation of SCD and SHD for millions of sequences can be time consuming and scales poorly with increasing chain length, making it less feasible for longer IDP designs. Although the Monte Carlo algorithm relies on the same shuffling and calculation methodologies, its motivated search was able to reach more significant extremes of LAF-1 RGG’s SCD sequence ensemble in a fraction of the iterations. As such, this tool provides an efficient means of designing polypeptide sequences that lie far outside the range of easily-accessible sequences.

**Fig 6.**
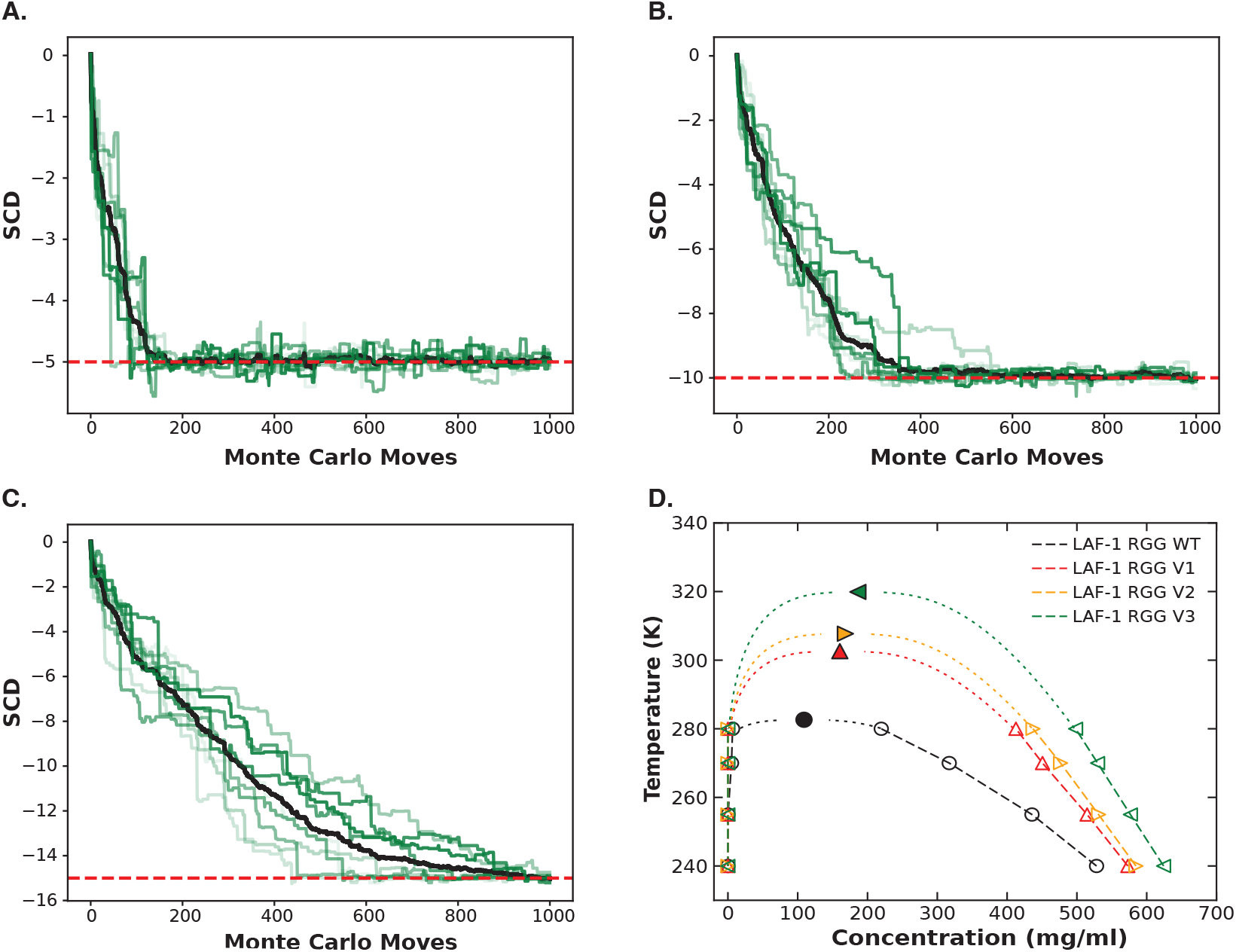
(A-C) 15 Monte Carlo runs starting with LAF-1 RGG and attempting to create variants with an SCD of −5 (A), −10 (B), and −15 (C). (D) Coexistence curve of LAF-1 RGG WT along with the 3 previously created variants.

We then tested these designed sequences using coarse-grained MD simulations in the HPS-Urry model with slab geometry [45]. The HPS-Urry model is well-suited for this problem as it has successfully been applied to capture single chain properties of a large library of IDPs, and also provide accurate phase diagram predictions for some IDPs with experimentally-measured phase diagrams [45]. We simulated the WT LAF-1 RGG as well as three designed variants with increasingly negative values of SCD, finding that more negative values of SCD resulted in upward shifts of the binodal phase envelope, indicating stronger self-association and greater phase separation propensity (Fig. 6D). We also determine the critical temperature for these variants (Table 1) and find that the observed increase in critical temperature is comparable to that reported for shuffled sequences designed in previous work [35]. The designed sequences are listed in S3 Table.

**Table 1.** SCD, SHD, SAD, and *T*_*c*_ values for LAF-1 RGG wild type and variants.

| Protein name | SCD | SHD | SAD | $T_c(K)$ |
| --- | --- | --- | --- | --- |
| LAF-1 RGG WT | 0.020 | 5.339 | 0.019 | 282.6 |
| LAF-1 V1 | -5.000 | 5.327 | 0.031 | 302.6 |
| LAF-1 V2 | -9.997 | 5.328 | 0.020 | 307.7 |
| LAF-1 V3 | -15.002 | 5.349 | 0.028 | 319.9 |

Beyond critical temperature, we also investigated whether this method of LLPS optimization influences the intrinsic material properties of the simulated condensate. We computed the single-chain translational diffusivity and the *R*_*g*_ autocorrelation time 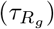 of LAF-1 RGG WT and variants V1–V3 across the temperatures used to construct the phase diagrams (Fig. 7). When plotted against temperature, all four sequences diffuse and re-orient faster at higher temperatures (Fig. 7A,C), with each designed variant diffusing and reconfiguring more slowly than the WT at the same temperature, and showing progressively slower dynamics for sequences with a greater degree of charge segregation. One major contribution to this effect could be the higher density of the more charge segregated variants in the dense phase. Indeed, when re-plotted against the dense-phase concentration (Fig. 7B,D), we find that the four sequences fall onto a much closer density-dependent trend, having lower diffusivity and longer 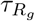 at higher dense-phase concentration. The designed sequences deviate only slightly from the WT, indicating that the density is indeed a major determinant of the materials properties of these dense phases. Importantly, slight variations between the different sequences at similar density cannot be explained by thermal effects because the designed variants are simulated at higher temperatures to achieve the same density as WT, but they exhibit slower dynamics and reconfiguration than the WT. Instead, this would likely be due to longer intrachain and interchain contact lifetimes [33].

**Fig 7.**
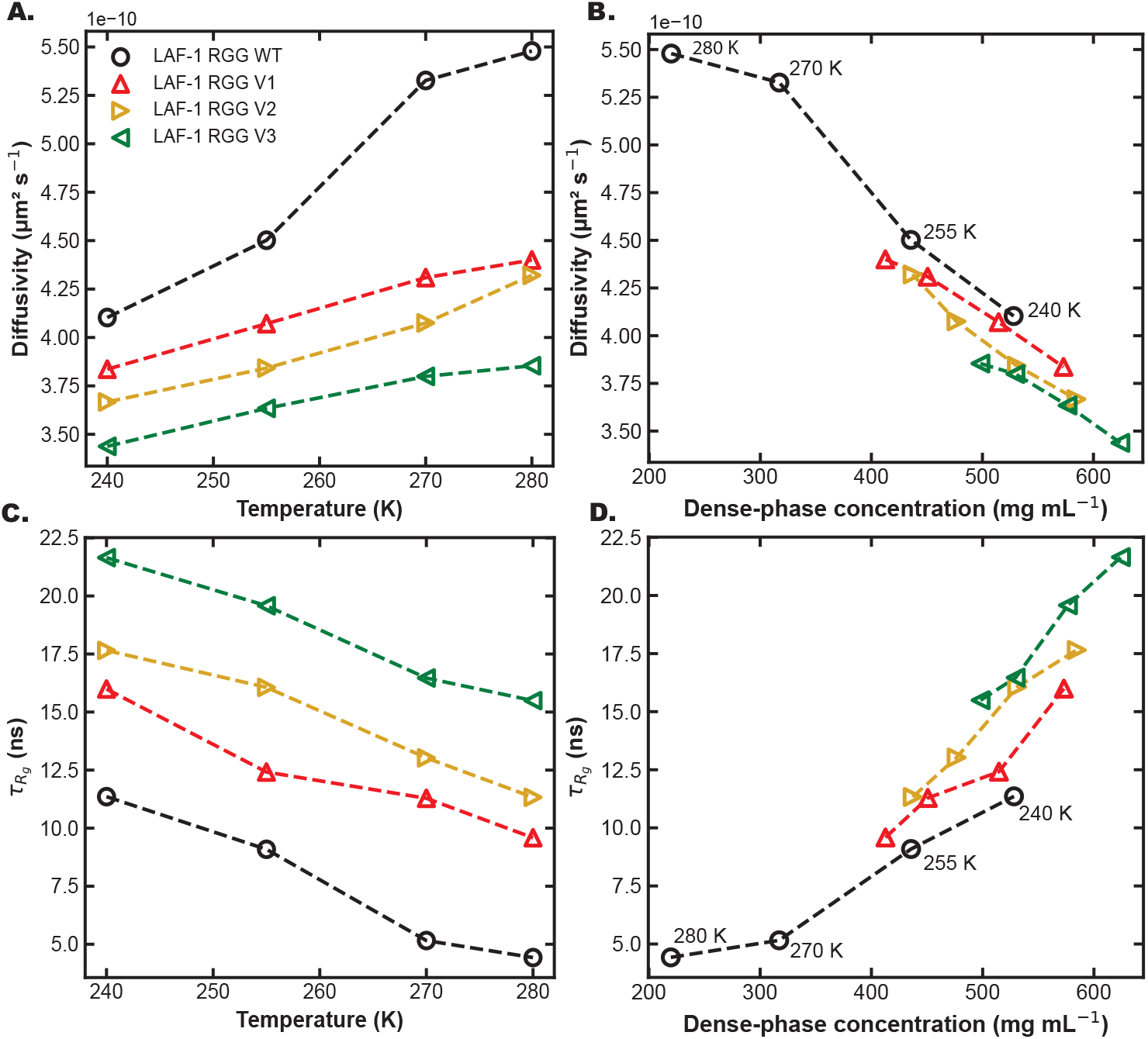
Effect of SCD-targeted optimization on intra-condensate dynamics of LAF-1 RGG. (A) Single-chain translational diffusivity in the dense phase versus temperature for LAF-1 RGG WT and the three designed variants V1, V2, V3 (Table 1). (B) The same diffusivity data plotted against the corresponding dense-phase concentration. (C) *R*_*g*_ autocorrelation time 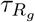 versus temperature for the same four sequences. (D) 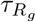 versus dense-phase concentration, with isotherms labeled.

We also investigated potential contributions of other parameters for influencing this increase in LLPS propensity for the designed variants. We find that *T*_*c*_ correlates most strongly with SCD (*r* = −0.97) and with Δ*G* (*r* = −0.96), but only weakly with SHD or SAD (S5 FigA). All of these results agree with previous studies showing that more negative SCD values correspond to more strongly segregated charged residues and such sequences exhibit higher LLPS propensity [21, 35, 36].

While SCD is the major driver of increased *T*_*c*_, we also wanted to determine if SHD could be designed to have a greater influence. Thus, we generated 15 additional fixed-composition LAF-1 RGG variants that nearly fully span the accessible range of SCD and SHD values (Fig. 8A; S4 Table). For each of these new variants, we conducted simulations and calculated the binodal phase diagram, and *T*_*c*_. Using this new dataset, we find that critical temperatures correlate strongly and monotonically with SCD (Fig. 8B; *r* = − 0.895), while showing essentially no dependence on SHD or SAD individually (S8 FigC,D). However, when we combine the three parameters in a multilinear regression model of *T*_*c*_, we find the agreement is moderately improved compared to predicting solely based on SCD (Fig. S8 FigA; r=-0.911). Our multilinear regression yields the standardized expression

**Fig 8.**
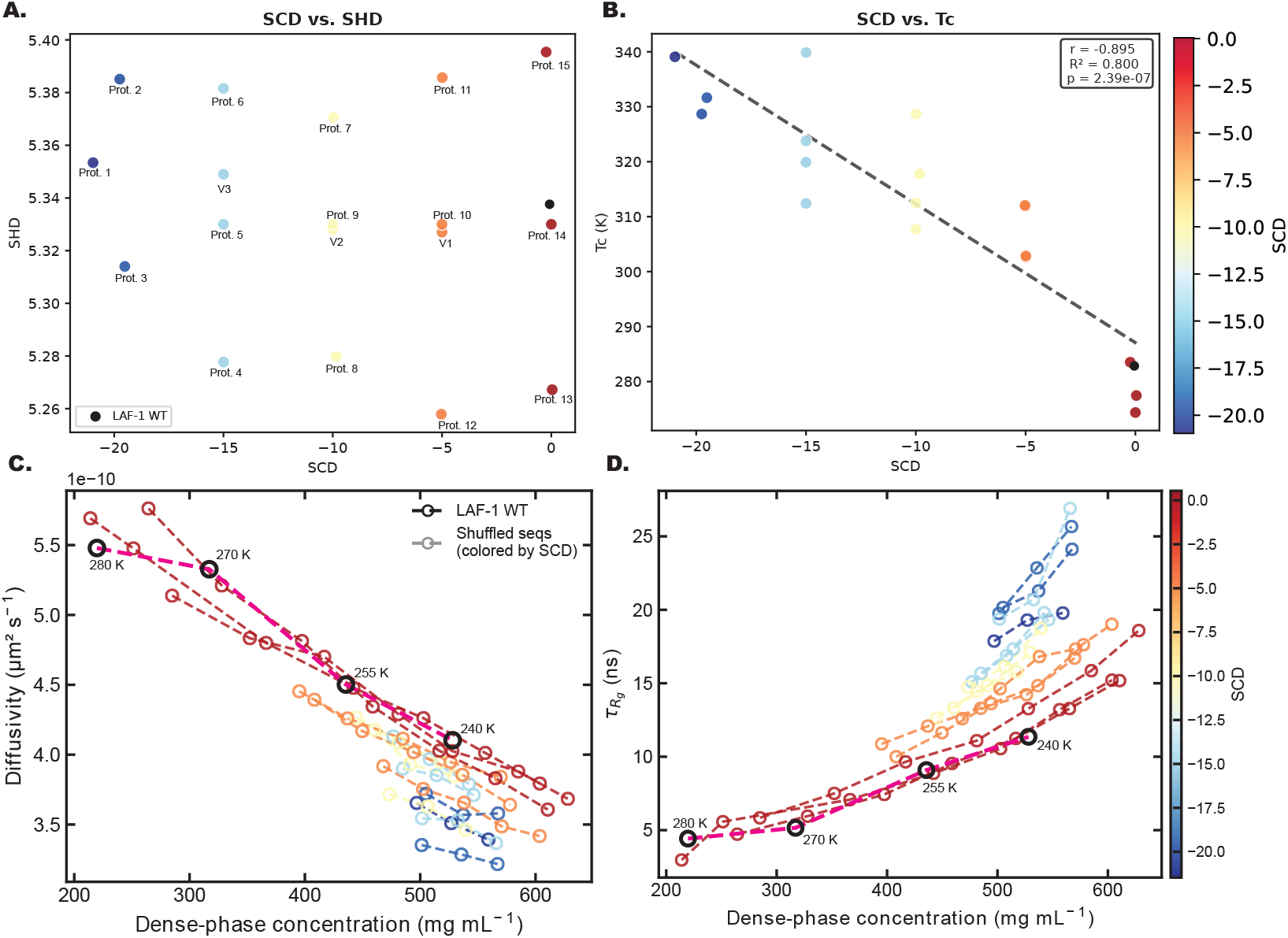
Fifteen additional fixed-composition LAF-1 RGG variants were designed to span a wide range of SCD and SHD values. (A) Coverage of the design library in the (SCD, SHD) plane, with WT shown as a black labeled marker. (B) Critical temperature Tc versus SCD. The strong, monotonic dependence of *T*_*c*_ on SCD, identifies SCD as the dominant determinant of LLPS stability across this composition-preserving library. (C) Single-chain diffusivity versus dense-phase concentration for the 15 designs at different temperatures, with points colored by SCD. (D) 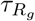versus dense-phase concentration, colored by SCD.

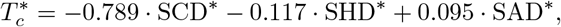

where each variable is *z*-score standardized (mean-centered and scaled by its standard deviation) so that coefficients are directly comparable. The associated feature importance ranking attributes ~ 79% of the explained variance in *T*_*c*_ to SCD, with SHD and SAD together contributing the remaining ~ 21% (S8 FigB). Taken together, these analyses indicate that, for sequences of fixed composition identical to that of LAF-1 RGG, the upward shift in *T*_*c*_ is predominantly controlled by enhanced charge segregation, and only minorly tuned by hydropathy patterning or aromatic distribution. However, it is likely that in sequences having fewer charged residues, the phase behavior would be less predominantly described by SCD, such as we find with our designs in the following sections (S5 Fig).

These 15 new designs also allow us to revisit the material properties question on a larger and more diverse set. When colored by SCD, the diffusivity-versus-density (Fig. 8C) and 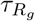-versus-density (Fig. 8D) curves overlap, indicating that dense-phase concentration, rather than SCD, is a significant determinant of the dynamics. The small differences that remain at a given density would not likely be due to temperature, since sequences with more negative SCD reach that density at a higher temperature, but generally exhibit slower dynamics as we saw previously with the smaller dataset (Fig. 6).

The same framework can be used in the opposite direction to generate variants with reduced phase-separation propensity relative to the WT. As an example, sequence protein14 in the extended LAF-1 RGG library has a *T*_*c*_ of 274.4 K (S4 Table), below the WT value of 282.6 K, and its coexistence curve lies below that of the WT (S7 Fig). This demonstrates that the Monte Carlo algorithm is not biased toward enhanced LLPS and can equally well design fixed-composition sequences that are less prone to phase separation. Combining all 18 LAF-1 RGG variants, we constructed an approximate 2D phase-behavior map in the (SCD, SHD) plane (S6 Fig), in which each sequence is annotated by its *T*_*c*_. The map reproduces the trend quantified above, *T*_*c*_ varies most strongly with SCD and only weakly with SHD at fixed composition, and provides a practical visual reference for the SCD*/*SHD ranges that demarcate strongly versus weakly phase-separating LAF-1 RGG variants in our model. We caution that the location of these boundaries is composition and force-field dependent, so for sequences whose composition departs substantially from LAF-1 RGG, the same (SCD, SHD) coordinates may correspond to a different phase regime.

### Adding new features to sequence optimization

Next, we incorporate an IDP’s predicted transfer free energy (Δ*G*) that directly quantifies the degree to which a sequence will phase separate, being related to the log of the IDP’s partition coefficient. To do this, we utilize the predictive algorithm from von Büllow et al. which predicts Δ*G* of phase separation directly from amino acid sequence [39]. This predictive algorithm was developed within an active-learning framework that iteratively combined coarse-grained molecular dynamics simulations with experimental data to train a neural network that maps amino acid sequence directly to the transfer free energy of phase separation (Δ*G*) and to the saturation concentration (c_*sat*_). The training set spans a diverse collection of intrinsically disordered regions (IDRs), and the predictor has been validated against both simulated and experimental LLPS data, including a proteome-wide application to the human IDRome [39]. We note that simulations used in training of this model were done using the CALVADOS 2 coarse-grained model [46] which employs a somewhat different hydropathy scale from the HPS-Urry scale used in this work. Thus our SHD parameter (and indeed the other patterning parameters) are likely to correlate somewhat with the ΔG predictions, as their influence on LLPS is likely to have been captured. Our intention in incorporating the Δ G predictions is to include a parameter that has some direct validation from experimental data to serve as a rough approximation of overall phase separation propensity, since none of our specific patterning parameters do a perfect job at predicting LLPS propensity in all cases (Fig. 8B; S5 Fig; S8 Fig).

We also introduce composition root-mean-square deviation (compRMSD) as a parameter that quantifies how much a designed sequence deviates from a reference sequence composition while allowing for Mutation moves.

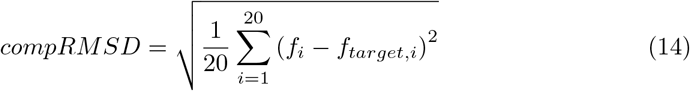

where f_*i*_ is the fraction of residues in the sequence that are amino acid i. Unlike fixed-composition approaches, enabling mutation moves and restraining overall composition drift through compRMSD enables controlled variation, making it particularly valuable for designing IDPs that retain similar composition to the template sequences, but may differ somewhat in size or total length. The weight of the compRMSD component can also be adjusted to modify the importance of preserving sequence composition. In the absence of mutation moves, FUS LC cannot achieve significantly lower Δ*G* values than the WT sequence, even with our optimization. With mutations enabled, it rapidly reaches the target value (S9 Fig).

By integrating these two parameters into our Monte Carlo sequence design approach, we designed two variants of the FUS LC wild-type (WT) protein: Variant 1 (V1) and Variant 2 (V2). As shown in the phase diagrams (Figure 9), both variants exhibit a lower Δ*G* compared to WT, correlating with an increased propensity for phase separation. These results demonstrate how incorporating additional predictive metrics, such as transfer free energy, allows for the rational design of IDP sequences with enhanced phase behavior. We find that, unlike with the LAF-1 RGG like sequences, the *T*_*c*_ for these FUS LC-derived sequences are predicted well from SHD and SAD, likely owing to the lower content of charged residues.

**Fig 9.**
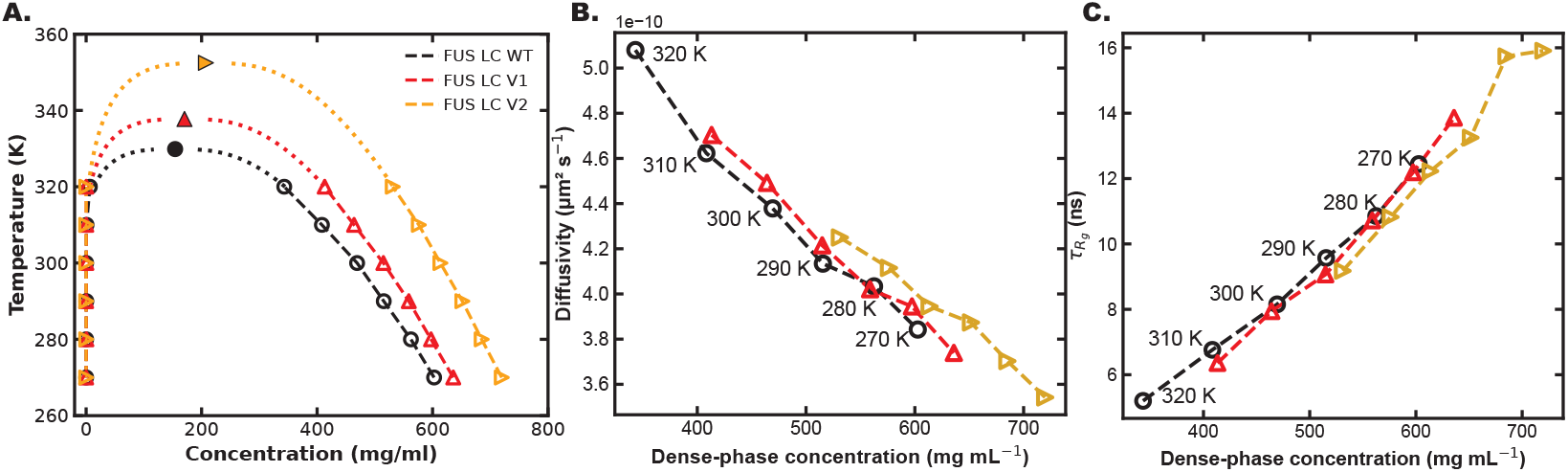
Phase behavior and intra-condensate dynamics of FUS LC WT and designed variants. (A) Temperature–concentration phase diagrams obtained from slab simulations for FUS LC WT and the two designed variants V1 and V2 (Table 2). Both variants exhibit upward-shifted binodals consistent with their more favorable predicted Δ*G*. (B) Single-chain translational diffusivity within the dense phase plotted against dense-phase concentration. (C) *R*_*g*_ autocorrelation time 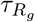 versus dense-phase concentration.

In summary, the designed FUS LC WT variants maintain compositions that are reasonably close to the wild type, with compRMSD values of 0.008 and 0.012 for variants V1 and V2, respectively (Table 2). By keeping compositional changes minimal, we ensure that the observed differences in predicted phase behavior primarily arise from altered residue patterning rather than shifts in overall amino acid content. The designed sequences are listed in S3 Table.

**Table 2.**
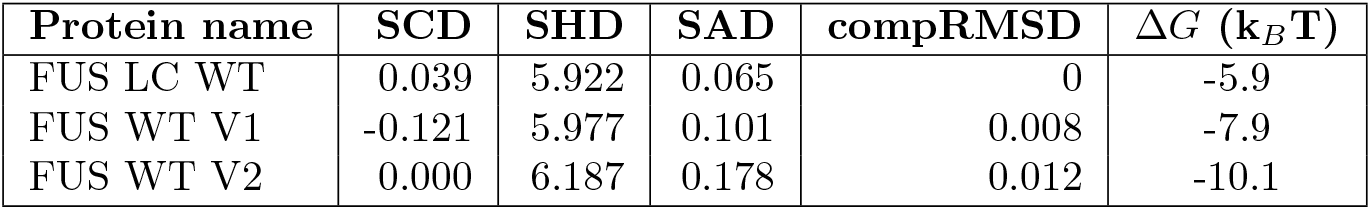
SCD, SHD, SAD, compRMSD, and ΔG values for FUS LC WT wild type and variants.

| Protein name | SCD | SHD | SAD | compRMSD | $\Delta G$ ($k_B T$) |
| --- | --- | --- | --- | --- | --- |
| FUS LC WT | 0.039 | 5.922 | 0.065 | 0 | -5.9 |
| FUS WT V1 | -0.121 | 5.977 | 0.101 | 0.008 | -7.9 |
| FUS WT V2 | 0.000 | 6.187 | 0.178 | 0.012 | -10.1 |

Beyond the critical temperature, we examined the intra-condensate dynamics of FUS LC WT and the two designed variants by computing single-chain translational diffusivity and the *R*_*g*_ autocorrelation time 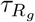. When these observables are plotted against temperature (S10 Fig), the variants are systematically slower than the WT at matched temperature, diffusivity decreases and 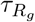 increases in the order WT →V1 →V2, mirroring their increasing phase-separation propensity. When the same data are instead plotted against dense-phase concentration (Fig. 9B,C), however, the three sequences fall onto a single common trend diffusivity decreases and 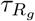 increases monotonically with dense-phase concentration. Notably, at matched dense-phase concentration the WT and the two variants are essentially indistinguishable unlike with the LAF-1 RGG derived variants. We note that because mutations were allowed (with compRMSD constraining but not eliminating compositional drift), the optimization raised the aromatic content from 24 residues in WT (14.7%) to 35 in V2 (21.5%), introducing 5 tryptophans and 2 phenylalanines absent from the WT and increased overall hydrophobic content from 24.5% to 31.3%, while the aromatic-decoration parameter SAD rose nearly threefold (0.065 →0.178; Table 2). Thus, the alterations to the sequence are more distributed throughout the length of the sequence and varied properties are changing.

Additionally, since FUS LC contains essentially no charged residues (≤3 in any variant), changes to aromatic and hydrophobic content and clustering is what drives the variants to phase separate at higher temperatures and to reach higher dense-phase concentrations. The minor variations in SCD are not enough to explain variations in phase separation propensity in this case (S5 FigB) providing a contrasting example to the previously designed LAF-1 RGG variants. In summary, we suggest that the total charge content is the major determinant of how much the phase behavior depends on SCD (S5 Fig).

### Leveraging design strategy for short peptides to mimic larger IDPs

It is well-established that shorter chain lengths, particularly in molecules with lower valence, are less prone to phase separation [28, 47–50]. However, recent work has shown that even short peptide sequences are capable of phase separation, usually owing to significant charge or aromatic content [27, 51, 52]. Short peptides have several advantages for experimental and computational research, including that simulations will converge more quickly and likely require fewer computing resources. We employed this method to design peptides that replicate the phase separation properties of FG Nucleoporin 153 (Nup153), a 1475-residue protein which is a component of the nuclear pore complex and implicated in binding to HIV-1 viral capsid proteins [53]. To demonstrate an extreme example of reducing protein sequence size, we have designed several miniature versions of Nup153, or mini-NUP sequences of length 30 (Table 3).

**Table 3.** SCD, SHD, SAD, compRMSD, and ΔG values for designed mini-NUP sequences. Values in parentheses are normalized values.

| Sequence | SCD | SHD | SAD | compRMSD | $\Delta G$ ($k_B T$) |
| --- | --- | --- | --- | --- | --- |
| mini-NUP seq1 | -0.80(-8.18) | 3.89(-1.98) | 0.23(-2.45) | 0.044 | -6.39 |
| mini-NUP seq2 | -0.72(-7.39) | 3.97(-2.05) | 0.32(-2.12) | 0.047 | -7.66 |
| mini-NUP seq3 | -0.25(-4.71) | 4.13(-3.66) | 0.32(-2.11) | 0.049 | -7.84 |

Since we have shortened the Nup153 sequence by a factor of 50, we expect that keeping identical composition will significantly reduce the phase separation propensity to a degree that it will not phase separate at accessible experimental conditions. Thus, we generated multiple variants of the peptide to have increasing likelihood of phase separation. To achieve the desired values of SHD, ΔG, and other sequence parameters, we increased the number of aromatic residues by introducing an additional 10% tyrosine and phenylalanine into the original FG Nucleoporin, Nup153. To quantify the compositional changes, we computed the compRMSD, which captures the difference in amino acid composition between the original sequence with added aromatic residues and the final designed sequences.

We conducted MD simulations of these peptide sequences to determine the phase separation propensity of each and find that we achieve a range of phase behaviors for each sequence (Fig. 10). The sequences of designed Nup153-like peptides are included in S3 Table.

**Fig 10.**
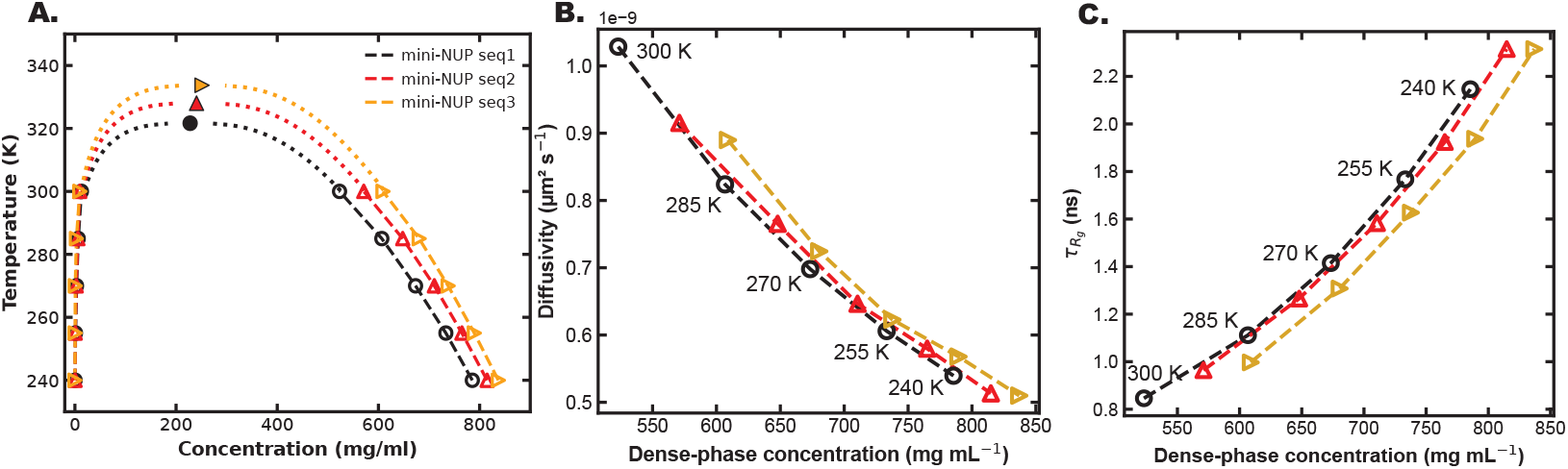
Phase behavior and intra-condensate dynamics of designed mini-NUP peptides. (A) Temperature–concentration phase diagrams obtained from slab simulations for the three 30-residue mini-NUP designs (`seq1`, `seq2`, `seq3`; Table 3). Despite all three peptides being only 30 residues long, the design framework yields variants that phase-separate across a broad range of conditions. (B) Single-chain translational diffusivity in the dense phase versus dense-phase concentration; isotherms (240, 255, 270, 285, 300 K) are labeled. (C) *R*_*g*_ autocorrelation time 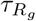versus dense-phase concentration, with the same isotherms labeled.

As with the LAF-1 RGG and FUS LC designs, we examined the intra-condensate dynamics of the three mini-NUP peptides by computing single-chain diffusivity and the *R*_*g*_ autocorrelation time 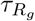 as a function of temperature (S11 Fig) and dense-phase concentration (Fig. 10B,C). One feature distinguishes these short peptides from the LAF-1 RGG case, *T*_*c*_ increases across the series seq1 →seq2 →seq3 (Table 3) while SCD becomes less negative (−0.80 →−0.72 →−0.25), the opposite of the trend established for LAF-1 RGG at fixed composition (Fig. 8B and S5 FigA), where *T*_*c*_ anticorrelates strongly with SCD. Inspection of the sequences (S3 Table) explains this reversal. In all three peptides the charges are strongly segregated—positively charged residues clustered near positions 3-6 and negatively charged residues near positions 22-27. The relevant difference between them is therefore not the patterning of charges but their total number, seq1 and seq2 each carry five charged residues, while seq3 carries only three. Because the design framework allowed mutations, the optimization satisfied the targeted increases in Δ*G* and SHD for seq3 by replacing two charged residues with aromatic/hydrophobic ones (aromatic fraction rises from 33.3% in seq1 to 36.7% in seq3, hydrophobic content from 50.0% to 60.0%, SAD rises from 0.23 to 0.32; Table 3). The less negative SCD in seq3 therefore does not indicate weaker charge patterning, the charges remain fully segregated, but rather fewer charges available to pattern, with the corresponding positions occupied by more cohesive residues. This compositional shift explains the reversed *T*_*c*_–vs–SCD trend, as the gain in aromatic/hydrophobic cohesion outweighs the small reduction in charge content. The intra-condensate dynamics, by contrast, behave just as they do for the LAF-1 RGG and FUS LC systems: when diffusivity and 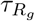 are plotted against temperature (S11 Fig), seq3 is the slowest and seq1 the fastest, mirroring their order of phase-separation propensity, but when the same observables are plotted against dense-phase concentration (Fig. 10B,C), the three peptides fall onto a single common trend and are essentially indistinguishable at matched dense-phase concentration and the small differences that remain may partly reflect temperature, since seq3 reaches a given dense-phase concentration at a higher temperature than seq1. As with the other systems, seq3 appears slower at matched temperature only because it reaches higher dense-phase concentrations, extending furthest along the common dynamics curve rather than occupying a distinct, more viscous material state. Taken together with the LAF-1 RGG and FUS LC results, this case illustrates two general points. First, intra-condensate dynamics are governed primarily by dense-phase concentration across all of our designs, so the optimization shifts the phase boundary without introducing a detectable change in material properties at matched density. Second, when mutations are enabled, SCD must be interpreted alongside composition, because it can change either through patterning at fixed composition (the LAF-1 RGG regime, where more negative SCD implies stronger LLPS) or through changes in charge content (the mini-NUP regime), only the former is captured by the fixed-composition heuristic.

## Methods

Simulations of designed IDP sequences were conducted using OpenMM simulation software [54] using the HPS-Urry coarse-grained model [45]. Each IDP was simulated at a range of temperatures where both a dense phase and a dilute phase were present to gather sufficient statistics to calculate the binodal coexistence curve. We ran simulations for at least 1 *µ*s per temperature case using Langevin dynamics. For all trajectories, the largest cluster was centered in the simulation box using an autocorrelation function of the instantaneous density profile of each simulation frame [55]. The critical point of each sequence was determined using a fitting function, assuming the universality of the 3D Ising model as in previous work [49].

### Force Field and Interaction Potentials

The HPS-Urry coarse-grained model was employed, where each amino acid is represented by a single bead located at the position of its *C*_*α*_ atom. The total interaction energy (*U*_total_) in this model is a sum of bonded and non-bonded terms:

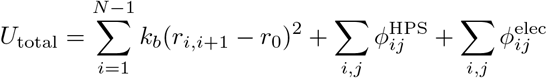

The first term models the bonded interactions between adjacent residues as a harmonic potential with a spring constant *k*_*b*_ = 20 kJ/Å^2^ and an equilibrium bond length of *r*_0_ = 3.82 Å. The non-bonded interactions are composed of hydrophobic 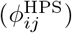 and electrostatic 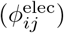 terms.

### Hydrophobic Interactions

These interactions are described by a modified Ashbaugh-Hatch potential [56], which is based on the Lennard-Jones (LJ) potential:

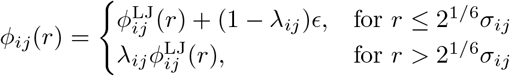

where the standard LJ potential is defined as:

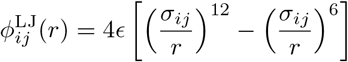

The energy scale is set by *ϵ* = 0.2 kcal/mol. The residue-specific parameters for size (*σ*_*i*_) and hydropathy/stickiness (*λ*_*i*_) are combined using an arithmetic mean: *σ*_*ij*_ = (*σ*_*i*_ + *σ*_*j*_)*/*2 and *λ*_*ij*_ = (*λ*_*i*_ + *λ*_*j*_)/2 - Δ, where Δ = 0.08.

### Electrostatic Interactions

Electrostatic forces are modeled using a Debye-Hückel potential to account for charge screening in solution [57]:

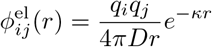

Here, *q*_*i*_ and *q*_*j*_ are the charges of the interacting residues. The dielectric constant of the solvent (*D*) was set to 80 to represent water, and the inverse Debye screening length (*κ*) was set to 1 nm^−1^, corresponding to a monovalent salt concentration of approximately 100 mM.

### Analysis: translational diffusivity and *R*_*g*_ relaxation time

Single-chain translational diffusivity (*D*) and the radius-of-gyration autocorrelation time 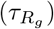 were computed from the same slab trajectories used to construct the binodal coexistence curves. All analysis was restricted to chains residing in the dense phase, identified as chains within the central high-density region of the slab. For the translational diffusivity, we tracked the position of the first C*α* bead of each chain and computed the mean-squared displacement (MSD) as a function of lag time *t*:

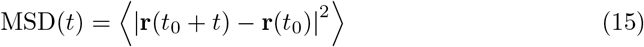

where the average is taken over all chains and all time origins *t*_0_, and displacements are evaluated under the minimum-image convention to account for periodic boundary conditions. Lag times up to 40 ns were computed (200 frames at a frame interval of 0.2 ns). The diffusion coefficient was extracted by fitting the MSD to the three-dimensional Einstein relation,

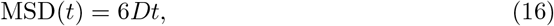

over the linear regime spanning 2–10 ns, giving *D* in units of *µ*m^2^ s^−1^. For the *R*_*g*_ autocorrelation time, the radius of gyration *R*_*g*_(*t*) was computed for each chain at every saved frame over the full production trajectory. The normalized autocorrelation function (ACF) was then calculated for each chain individually using a fast Fourier transform (FFT) algorithm:

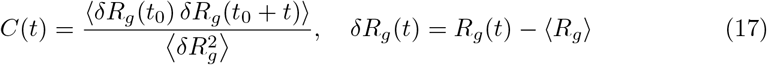

and averaged over all chains in the simulation. Lag times spanning the full production trajectory (up to ~1 *µ*s) were used. The relaxation time 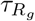 was extracted by fitting the chain-averaged ACF to a single exponential,

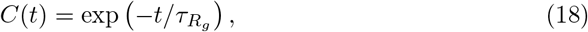

restricted to the region where *C*(*t*) ≥0.05 to avoid the noisy long-lag tail. All trajectory analysis was performed using `MDAnalysis` and `MDTraj`.

## Conclusion

By defining trends between composition-specific terms and patterning parameters, we have successfully established a method of approximating the properties of a given IDP’s constant-composition ensemble, and thus derived a normalization scheme for the common patterning parameters, SCD and SHD. Using this normalization scheme, we then leverage this into a method of comparing changes between different patterning parameters, putting them on an equal playing field in the design space. We test this out with an IDP sequence design Monte Carlo algorithm which can simulataneously optimize multiple design critera. Existing literature linking patterning parameters and LLPS further allows this Monte Carlo to represent a means of tuning the phase behavior of a given IDP by creating variants. We first demonstrate that our code can design extreme outliers of specific patterning parameters such as super charge-segregated LAF-1 RGG variants and, using an extended library of 15 additional fixed-composition LAF-1 RGG designs together with a multilinear regression of Tc on standardized (SCD, SHD, SAD), we show that for this charge-rich system the critical temperature is overwhelmingly controlled by charge patterning (Fig. 8, S8 Fig). We apply this method to design variants of FUS that are more prone to LLPS than the wild-type, focusing primarily on predicted ΔG of phase separation. We also design mini-NUP variants that maintain similar composition to the base sequence, Nup153, while scaling the importance of perfect compositional similarity to enable design of more “sticky” sequences more capable of phase separation, offsetting the significantly reduced size of the sequence.

Beyond critical temperatures, we further examined intra-condensate dynamics for each designed system and find that the LAF-1 RGG variants preserve the WT mobility density relationship at matched dense-phase concentration, whereas the FUS LC and mini-NUP variants for which mutations were enabled and composition was allowed to drift exhibit lower diffusivity and longer 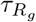 at matched density. This indicates that the same framework can be used either to shift the phase boundary alone or to jointly tune the phase boundary and the material properties of the resulting condensate.

Looking forward, this work can be supplemented by inclusion of multiple other parameters for optimization, such as: secondary structure predictors, disorder predictors or any other sequence-based parameter. Furthermore, as the link between patterning parameters of an IDP and the properties of its resultant condensate continue to be elucidated, this work could incorporate condensate properties more directly into its loss function rather than only the general propensity for phase separation. For example, as SAD seems to indicate a more gel-like condensate, this work could be improved to give users better control over designed condensate properties by giving additional guidance into what parameters of SAD may correlate with liquid-like and gel-like condensates. The dynamics analyses introduced for the FUS LC and mini-NUP designs in this work are a first step in that direction, and we anticipate that pairing them with the experimental measurements outlined above will be the most effective route to validating and refining these material-property aware design criteria. Since the present study is entirely computational, we present the designed variants as testable hypotheses. Finally, we emphasize that design of fully shuffled or randomized IDP sequences could impose other unexpected changes to the sequence behavior, perhaps beyond what can be predicted by our methodology. Thus it is important to have a good understanding of the base protein sequence, its overall composition and patterning properties as well as any important motifs or domains that might be influential to its function before attempting to make drastic perturbations to the sequence.

## Data and Code Availability Statement

All data used to generate figures in this work will has been deposited on an online database which can be accessed at https://zenodo.org/records/20397905. Source code for Monte Carlo sequence design software is also available on the same link, and on Github https://github.com/Dignon-Lab/Monte-Carlo. Additional data such as raw simulation trajectories will be available upon reasonable request.

## Supporting information

**S1 Fig**. Correlation of different predictive parameters with ensemble-calculated mean and standard deviation of SCD.

**S2 Fig**. Correlations of the patterning terms SCD, SHD, and SAD for 10^5^ shuffles of the IDPs FUS LC, LAF-1 RGG, and NUP-153.

**S3 Fig**. FUS LC Monte Carlo runs with no mutations. Multiple parameters are tracked but the trajectories only optimize for targets of the maximum (blue) and minimum (green) possible SCD values.

**S4 Fig**. Energy decomposition of five parameters with respect to mutations, swaps, and shuffles separately. These also include unaccepted Monte Carlo moves. The range of values observed for the three patterning parameters are within an a few-fold difference for each move type, indicating success in the normalization scheme for application to MC sequence design.

**S5 Fig**. Per-system correlations between critical temperature and sequence patterning metrics for the designed variants.(A) LAF-1 RGG variants *T*_*c*_ plotted against SCD, SHD, SAD, and the predicted Δ*G*. Pearson correlation coefficients are indicated in each panel. (B) FUS LC variants. (C) The three mini-NUP designs.

**S6 Fig**. Approximate 2D phase-behavior map for fixed-composition LAF-1 RGG designs. All 18 LAF-1 RGG variants (WT plus V1–V3 from Table 1 and the 15 additional designs in Table S4) are placed at their (SCD, SHD) coordinates and annotated by their simulated critical temperature.

**S7 Fig**. LAF-1 RGG variant with attenuated phase separation. Coexistence curves obtained from slab simulations for LAF-1 RGG WT (*T*_*c*_ = 282.6 K) and `protein14` (Table S4; *T*_*c*_ = 274.4 K), a fixed-composition variant designed by the same Monte Carlo procedure.

**S8 Fig**. Multilinear regression of critical temperature on patterning features for LAF-1 RGG designs. (A) Predicted versus observed *T*_*c*_ across the 18 LAF-1 RGG variants from a multilinear regression on the *z*-score standardized features. (B) Relative feature importance derived from the standardized coefficients. (C) Tc versus SHD. (D) *T*_*c*_ versus SAD.

**S9 Fig**. FUS LC Monte Carlo runs optimizing for only ΔG with (A) no mutations and (B) mutations enabled.

**S10 Fig**. Temperature dependence of intra-condensate dynamics for FUS LC variants. (A) Single-chain translational diffusivity in the dense phase versus temperature for FUS LC WT, V1, and V2. (B) *R*_*g*_ autocorrelation time 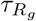 versus temperature for the same three sequences.

**S11 Fig**. Temperature dependence of intra-condensate dynamics for the mini-NUP designs. (A) Single-chain translational diffusivity within the dense phase versus temperature for mini-NUP `seq1`, `seq2`, and `seq3`. (B) *R*_*g*_ autocorrelation time 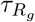 versus temperature for the same three sequences.

**S1 Table**. Urry normalized hydropathy values used for each amino acid.

**S2 Table**. List of amino acid sequences for real IDPs used in initial shuffling study.

**S3 Table**. List of designed sequences used in this work in MD simulations of phase separation.

**S4 Table**. List of 15 additional designed LAF-1 RGG variants generated to span the (SCD, SHD) design space at fixed amino acid composition.

**S1 Appendix**. Supporting information document with 11 figures, and 4 tables.

## Acknowledgments

G.L.D acknowledges support from NIH Grant R35GM150589 and start-up funds from Rutgers University, and computing resources through NSF ACCESS DISCOVER project CHM250012. We thank José Villegas and Ben Schuster for useful discussions.

## Supporting Information

### SI Figures

**Figure 1:**
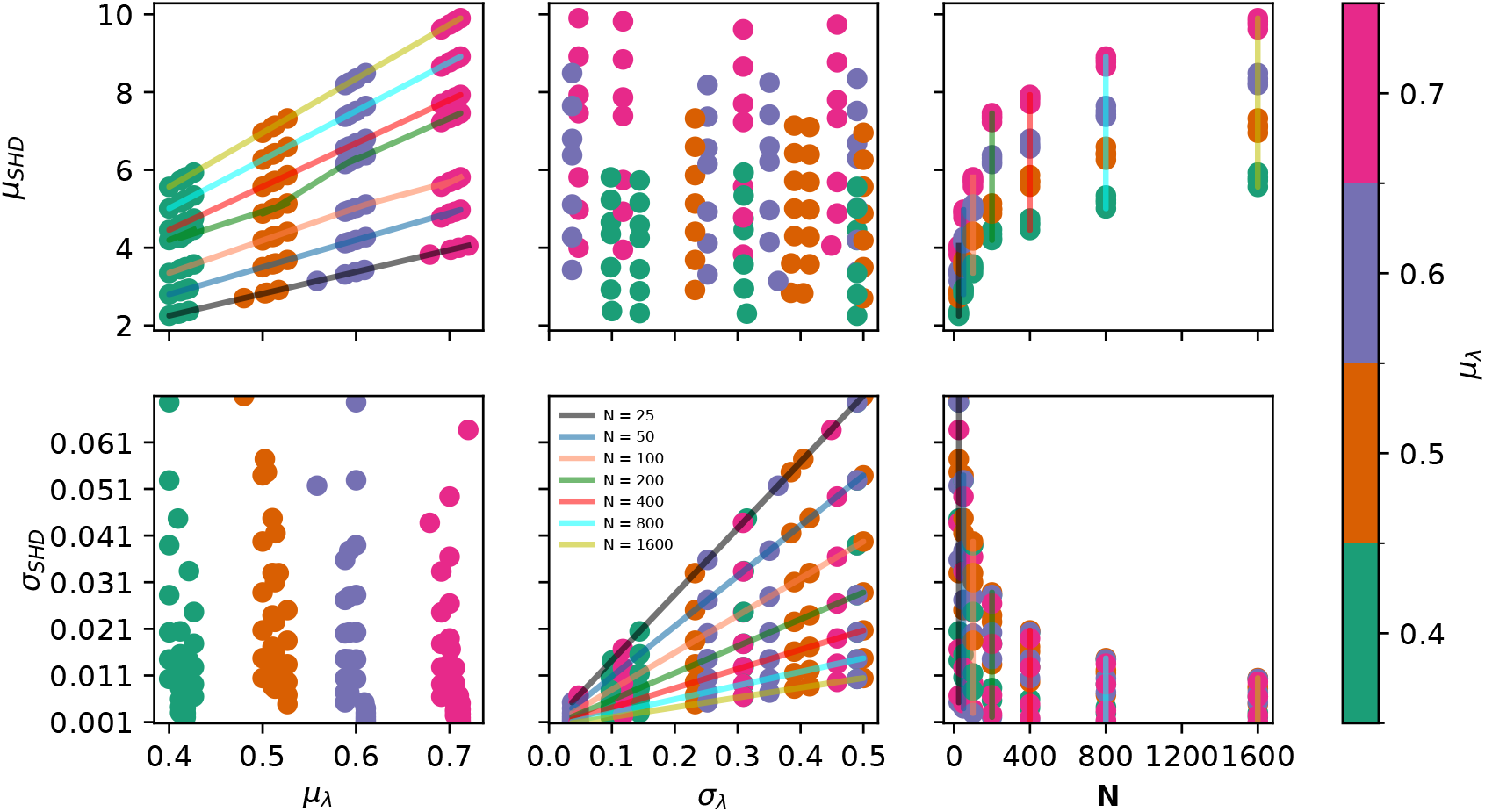
Correlation of different predictive parameters with ensemble-calculated mean and standard deviation of SHD. Lines connect points of the same length to show trends, they do not represent empirical fits.

**Figure 2:**
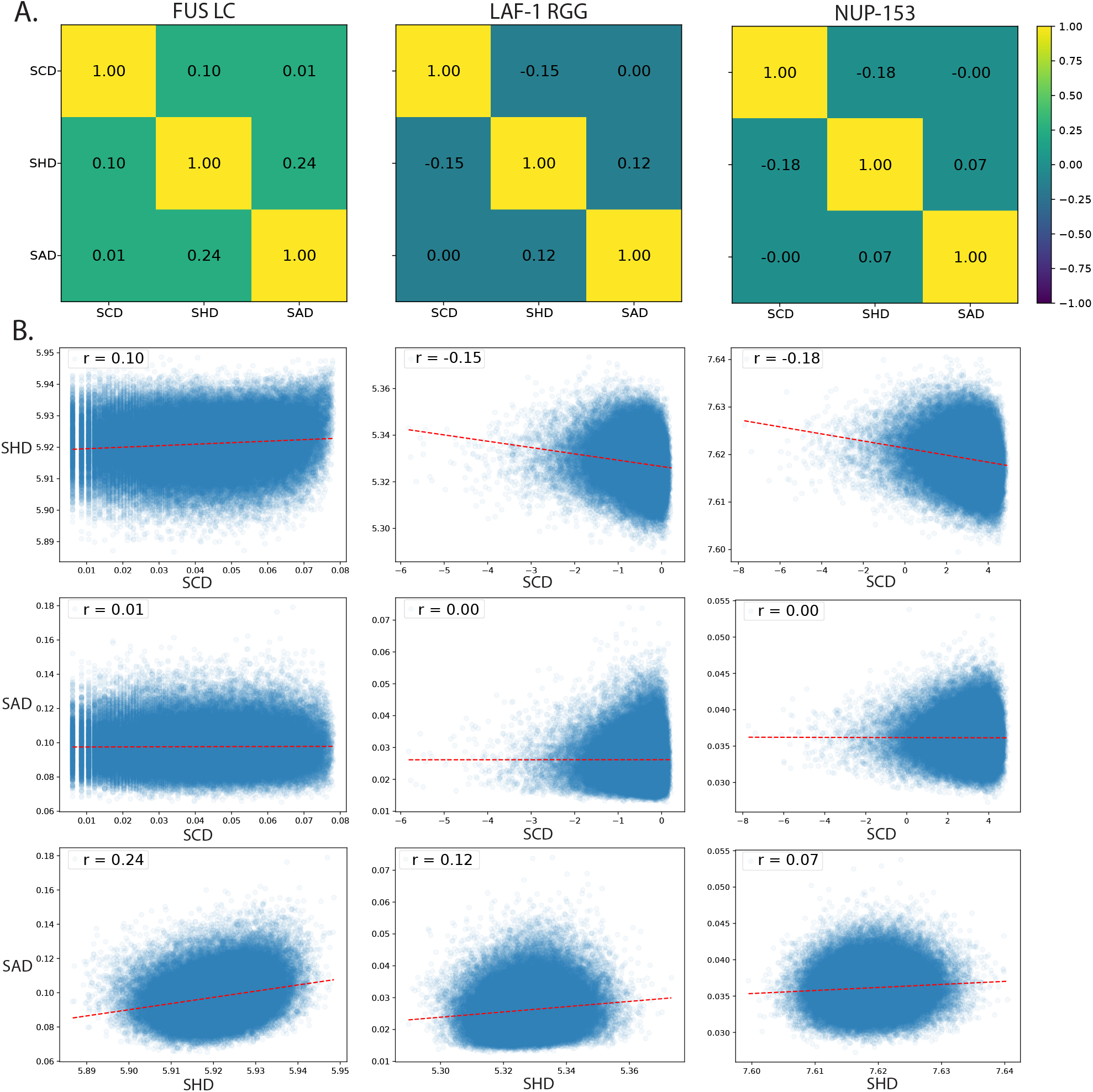
Correlations of the patterning terms SCD, SHD, and SAD for 10^5^ shuffles of the IDPs FUS LC, LAF-1 RGG, and NUP-153.

**Figure 3:**
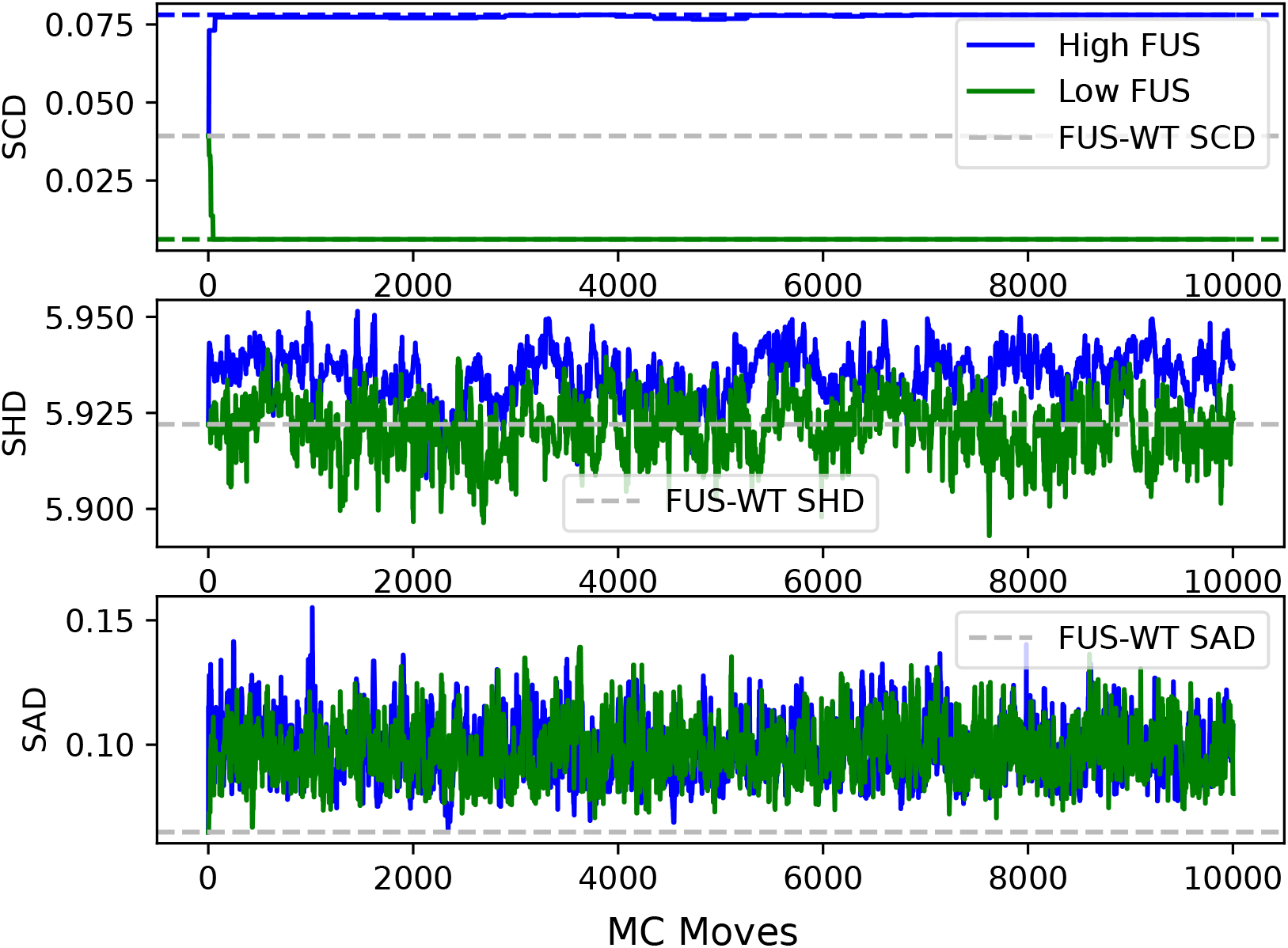
FUS LC Monte Carlo runs with no mutations. Multiple parameters are tracked but the trajectories only optimize for targets of the maximum (blue) and minimum (green) possible SCD values.

**Figure 4:**
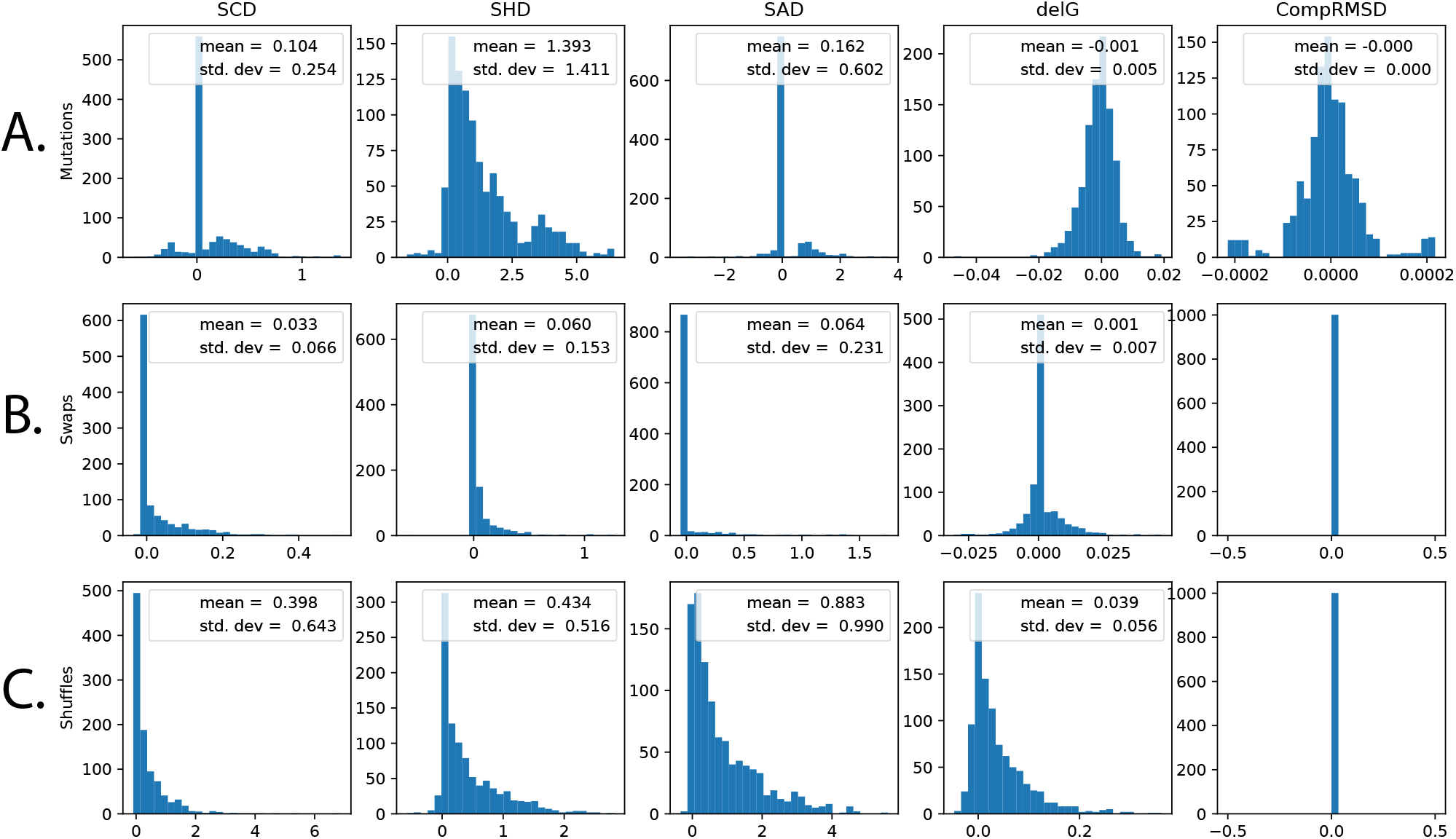
Energy decomposition of five parameters with respect to mutations, swaps, and shuffles separately. These also include unaccepted Monte Carlo moves. The range of values observed for the three patterning parameters are within an a few-fold difference for each move type, indicating success in the normalization scheme for application to MC sequence design.

**Figure 5:**
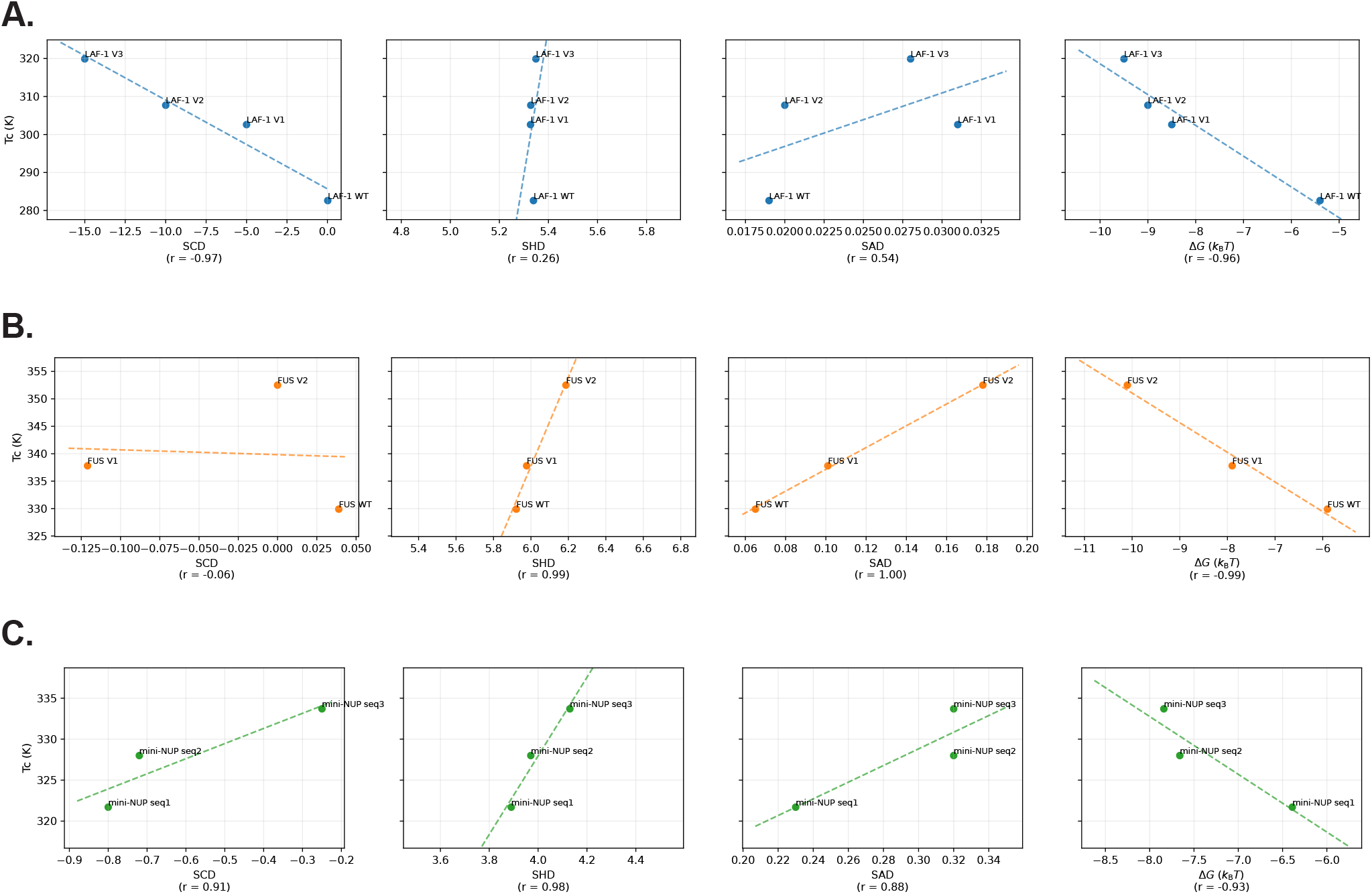
Per-system correlations between critical temperature and sequence patterning metrics for the designed variants. (A) LAF-1 RGG variants *T*_*c*_ plotted against SCD, SHD, SAD, and the predicted Δ*G*. Pearson correlation coefficients are indicated in each panel. (B) FUS LC variants. (C) The three mini-NUP designs.

**Figure 6:**
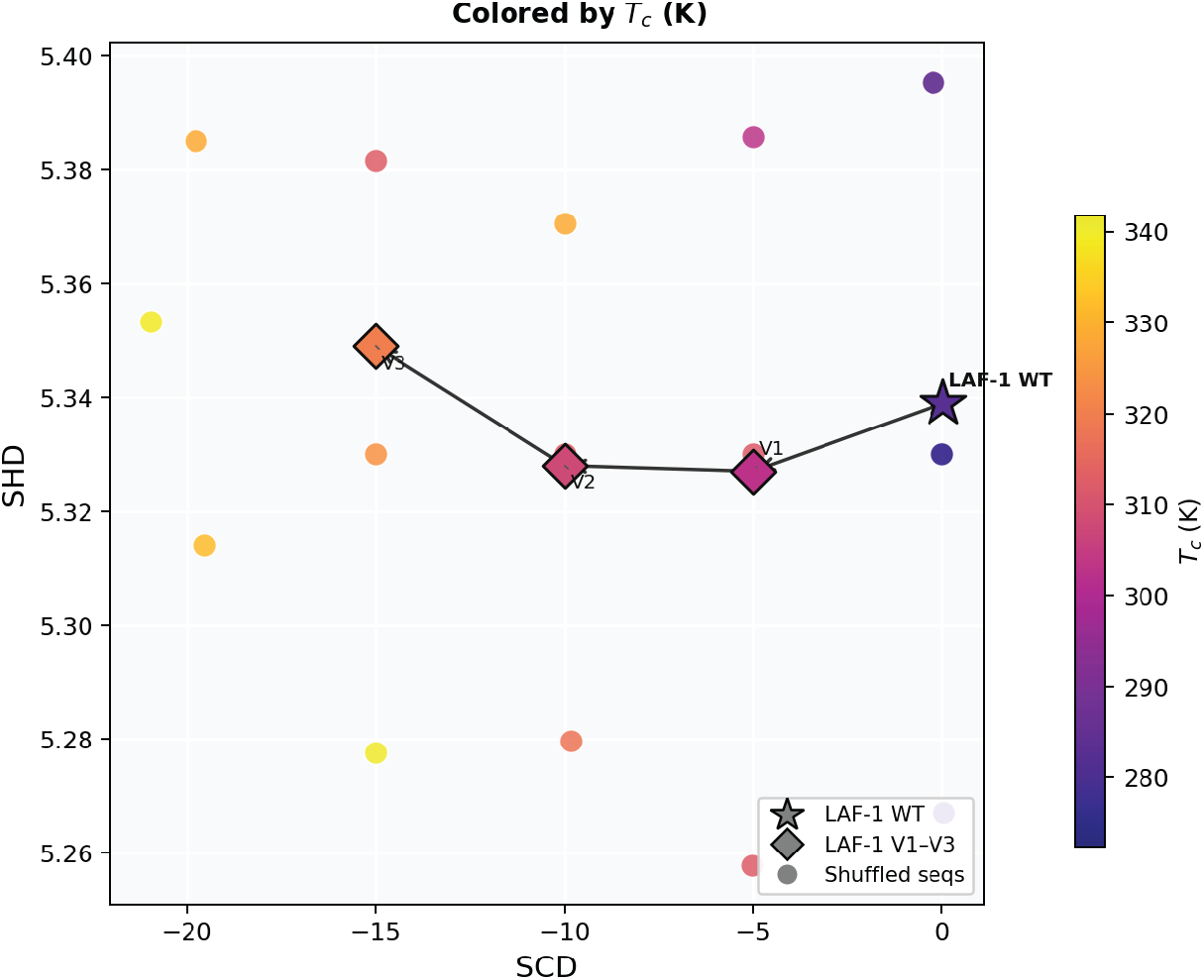
Approximate 2D phase-behavior map for fixed-composition LAF-1 RGG designs. All 18 LAF-1 RGG variants (WT plus V1–V3 from Table 1 and the 15 additional designs in Table S4) are placed at their (SCD, SHD) coordinates and annotated by their simulated critical temperature.

**Figure 7:**
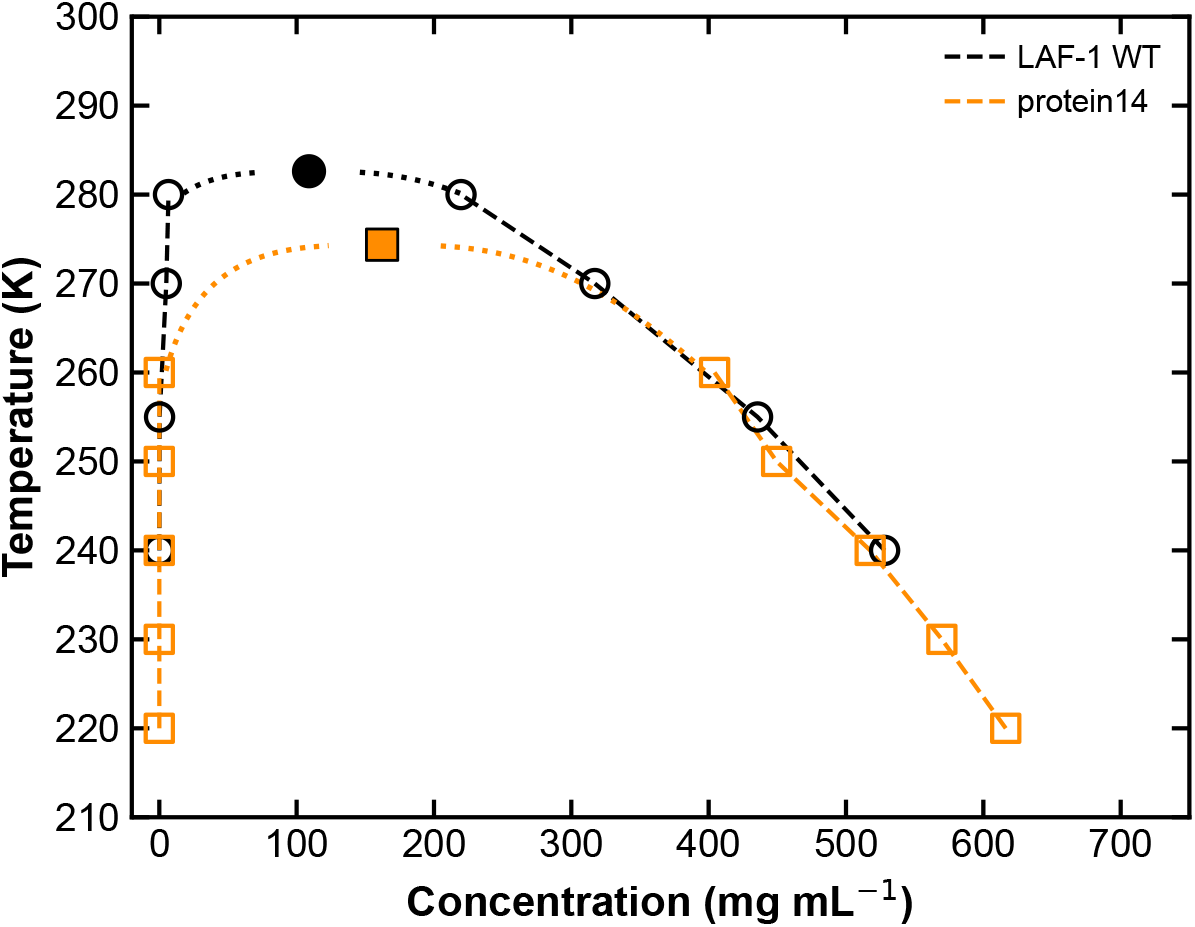
LAF-1 RGG variant with attenuated phase separation. Coexistence (binodal) curves obtained from slab simulations for LAF-1 RGG WT (*T*_*c*_ = 282.6 K) and `protein14` (Table S4; *T*_*c*_ = 274.4 K), a fixed-composition variant designed by the same Monte Carlo procedure.

**Figure 8:**
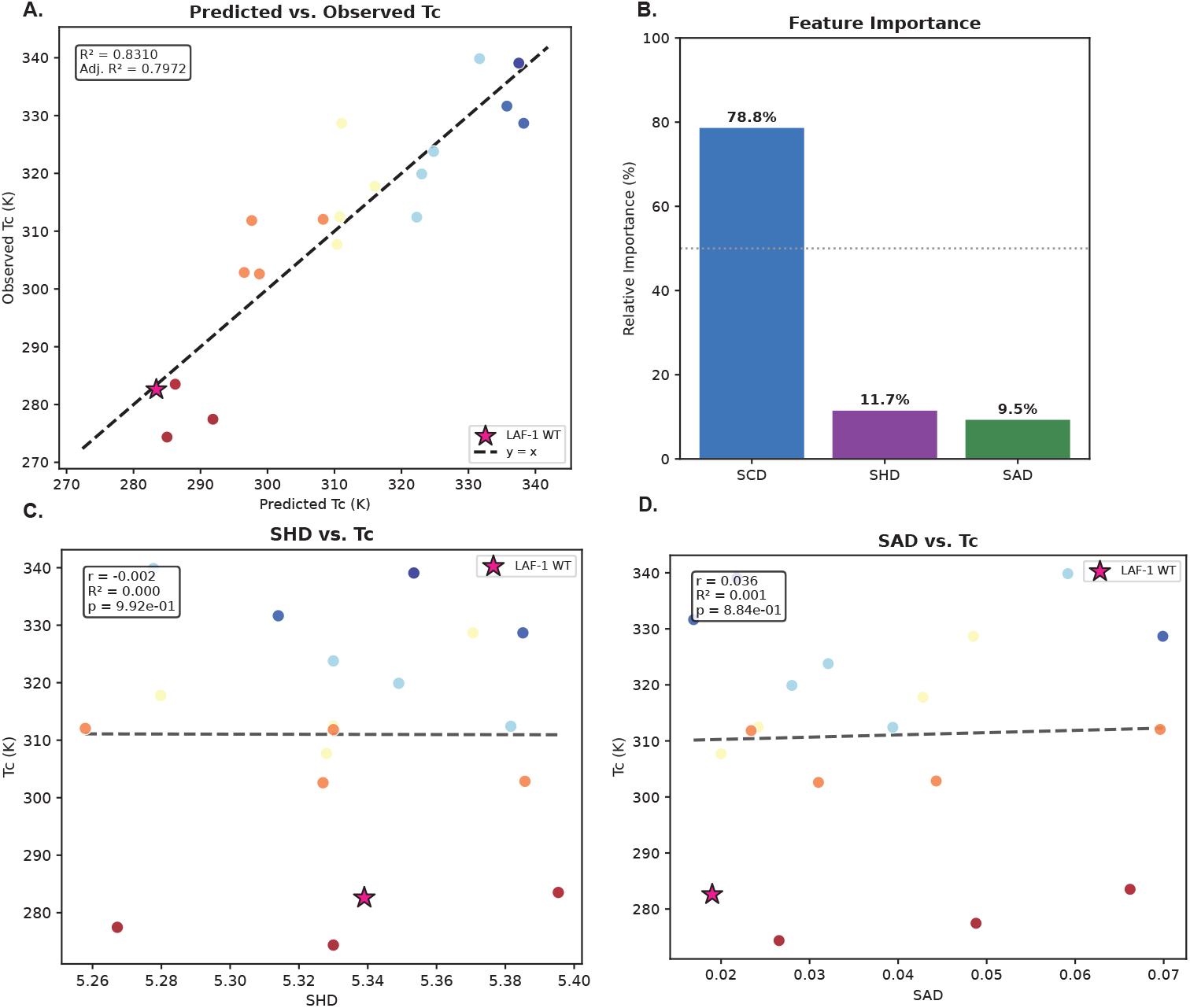
Multilinear regression of critical temperature on patterning features for LAF-1 RGG designs. (A) Predicted versus observed *T*_*c*_ across the 18 LAF-1 RGG variants from a multilinear regression on the *z*-score standardized features. (B) Relative feature importance derived from the standardized coefficients. (C) Tc versus SHD. (D) *T*_*c*_ versus SAD.

**Figure 9:**
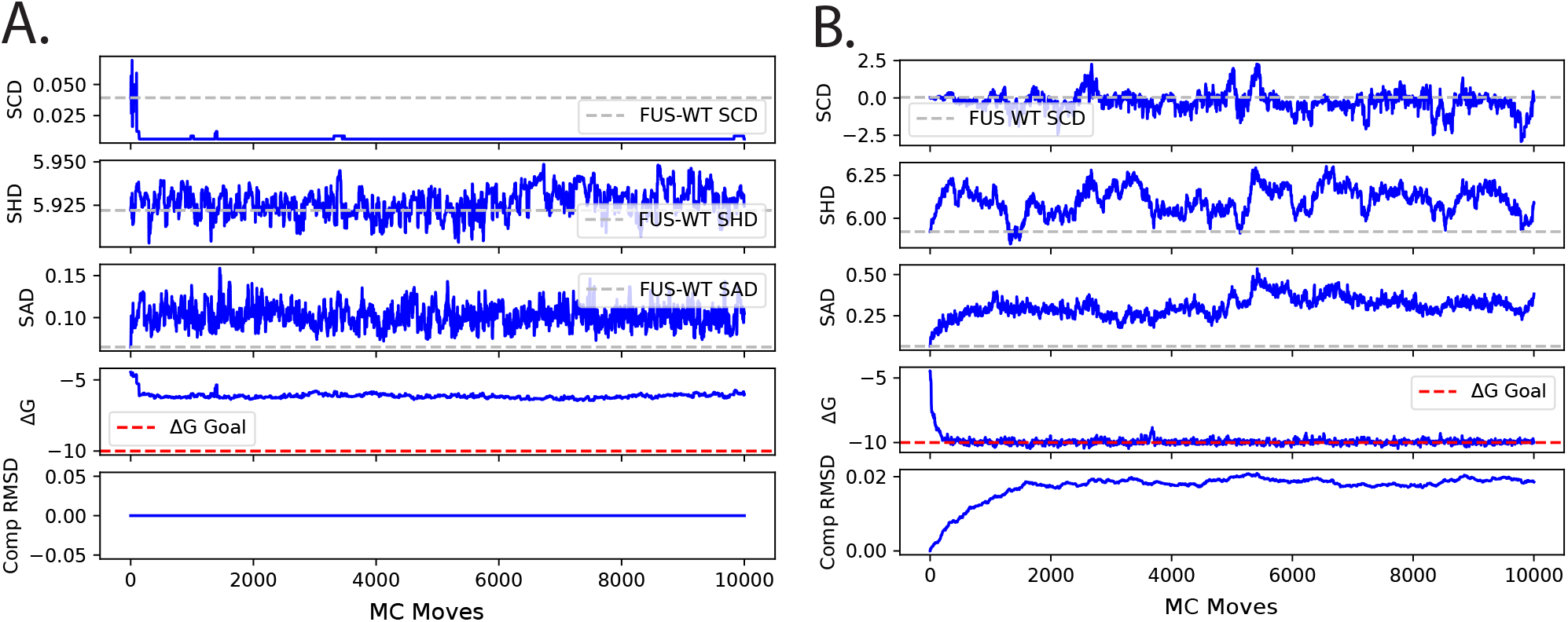
FUS LC Monte Carlo runs optimizing for only ΔG with (A) no mutations and (B) with mutations enabled.

**Figure 10:**
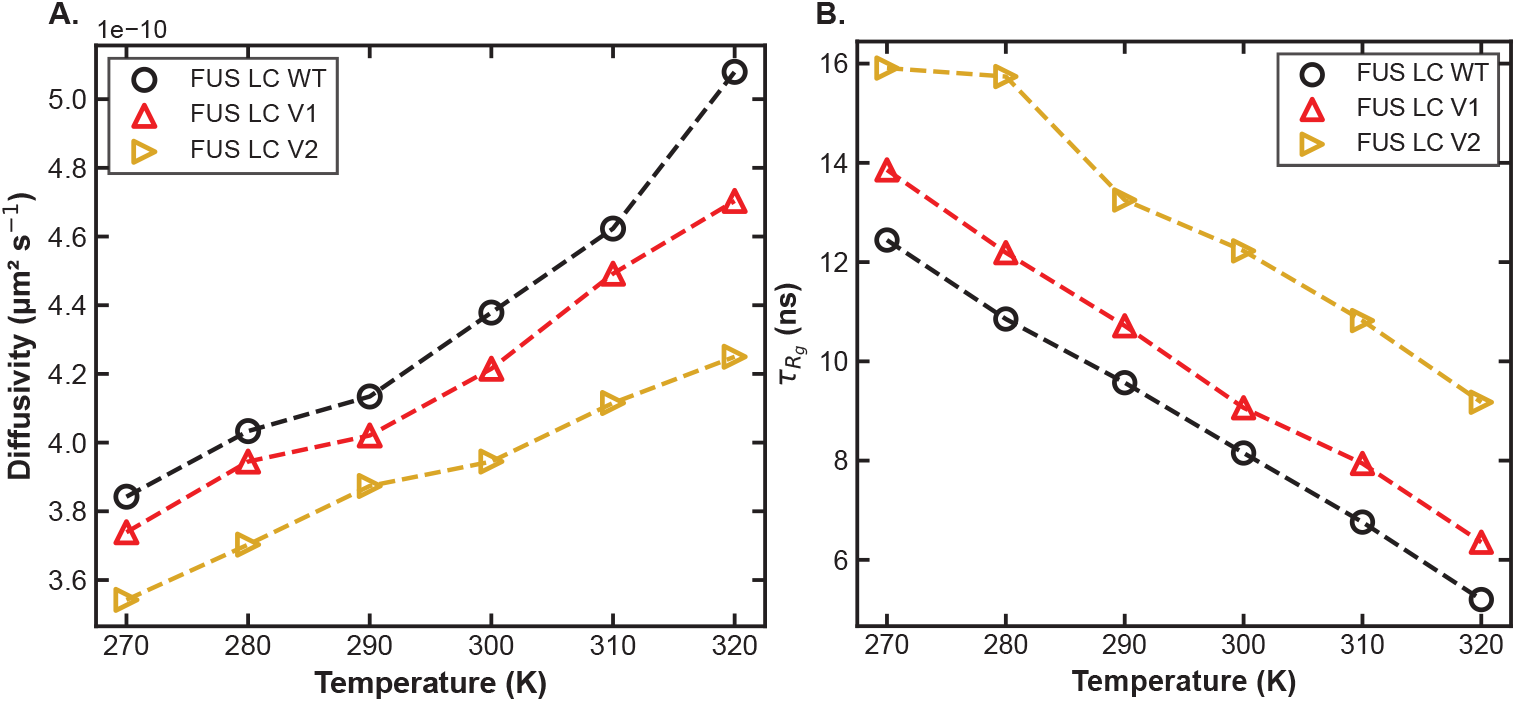
Temperature dependence of intra-condensate dynamics for FUS LC variants. (A) Single-chain translational diffusivity in the dense phase versus temperature for FUS LC WT, V1, and V2. (B) *R*_*g*_ autocorrelation time 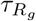 versus temperature for the same three sequences.

**Figure 11:**
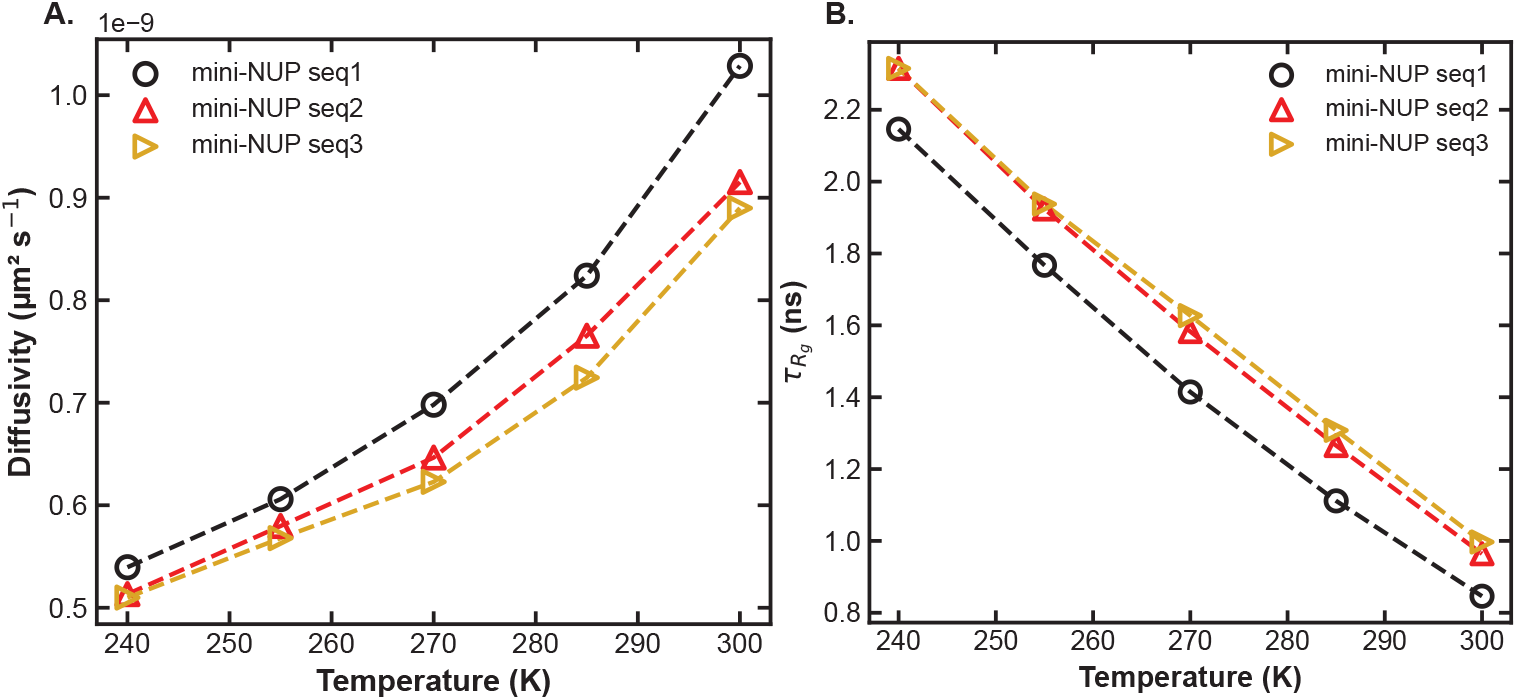
Temperature dependence of intra-condensate dynamics for the mini-NUP designs. (A) Single-chain translational diffusivity within the dense phase versus temperature for mini-NUP `seq1`, `seq2`, and `seq3`. (B) *R*_*g*_ autocorrelation time 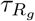 versus temperature for the same three sequences.

### SI Tables

**Table 1:** Normalized hydropathy scales used for the HPS-Urry models.

| Amino acid | Urry [1] normalized hydrophathy scale ($\lambda$) |
| --- | --- |
| TRP | 1.000000 |
| TYR | 0.897059 |
| PHE | 0.823530 |
| HIS | 0.764707 |
| PRO | 0.758824 |
| LEU | 0.720589 |
| ILE | 0.705883 |
| MET | 0.676471 |
| VAL | 0.664707 |
| CYS | 0.647060 |
| ALA | 0.602942 |
| ASN | 0.588236 |
| SER | 0.588236 |
| THR | 0.588236 |
| GLY | 0.573530 |
| ARG | 0.558824 |
| GLN | 0.558824 |
| LYS | 0.382354 |
| ASP | 0.294119 |
| GLU | 0.000000 |

**Table 2:**
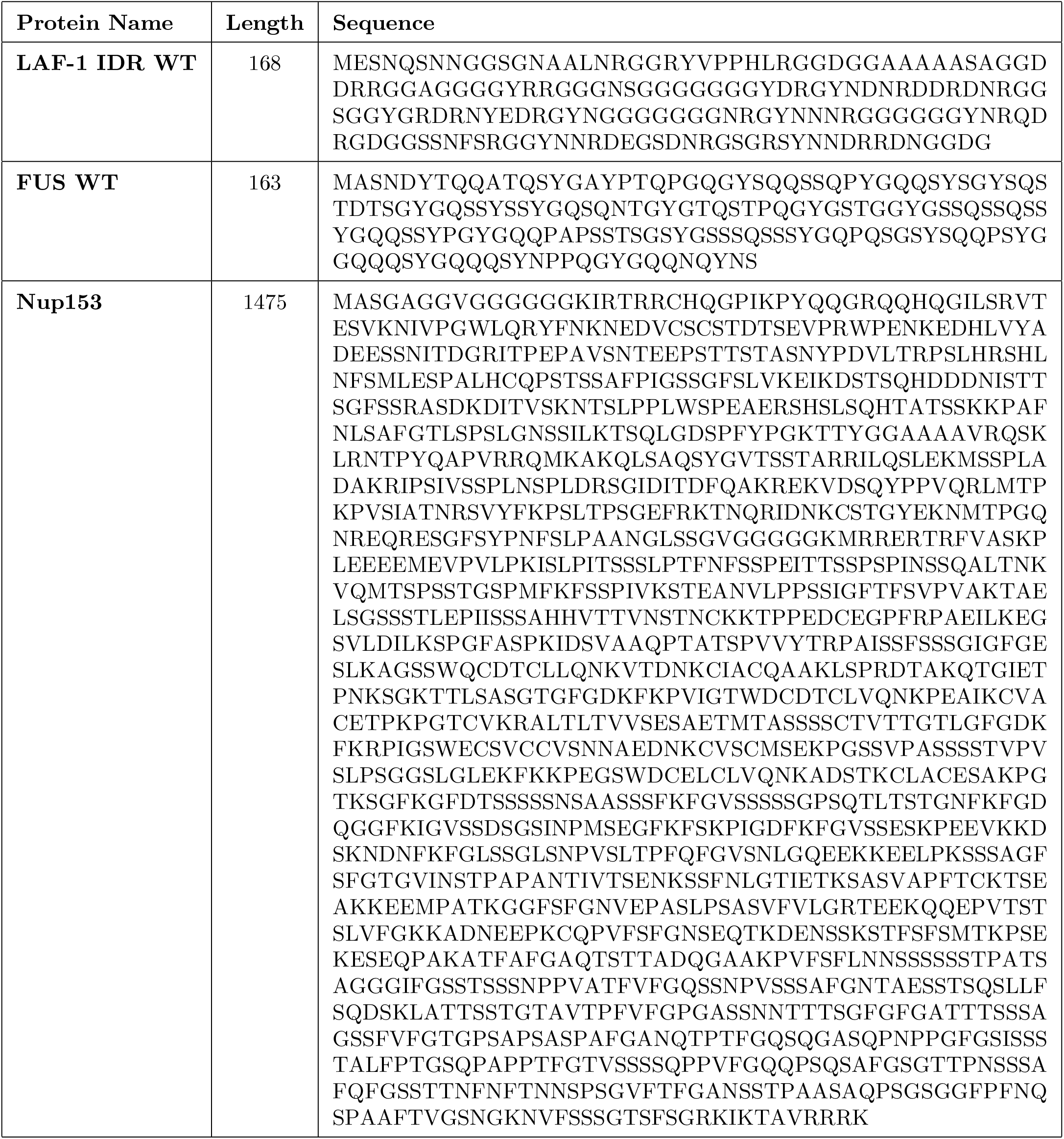
List of amino acid sequences for real IDPs used in initial shuffling study.

**Table 3:**
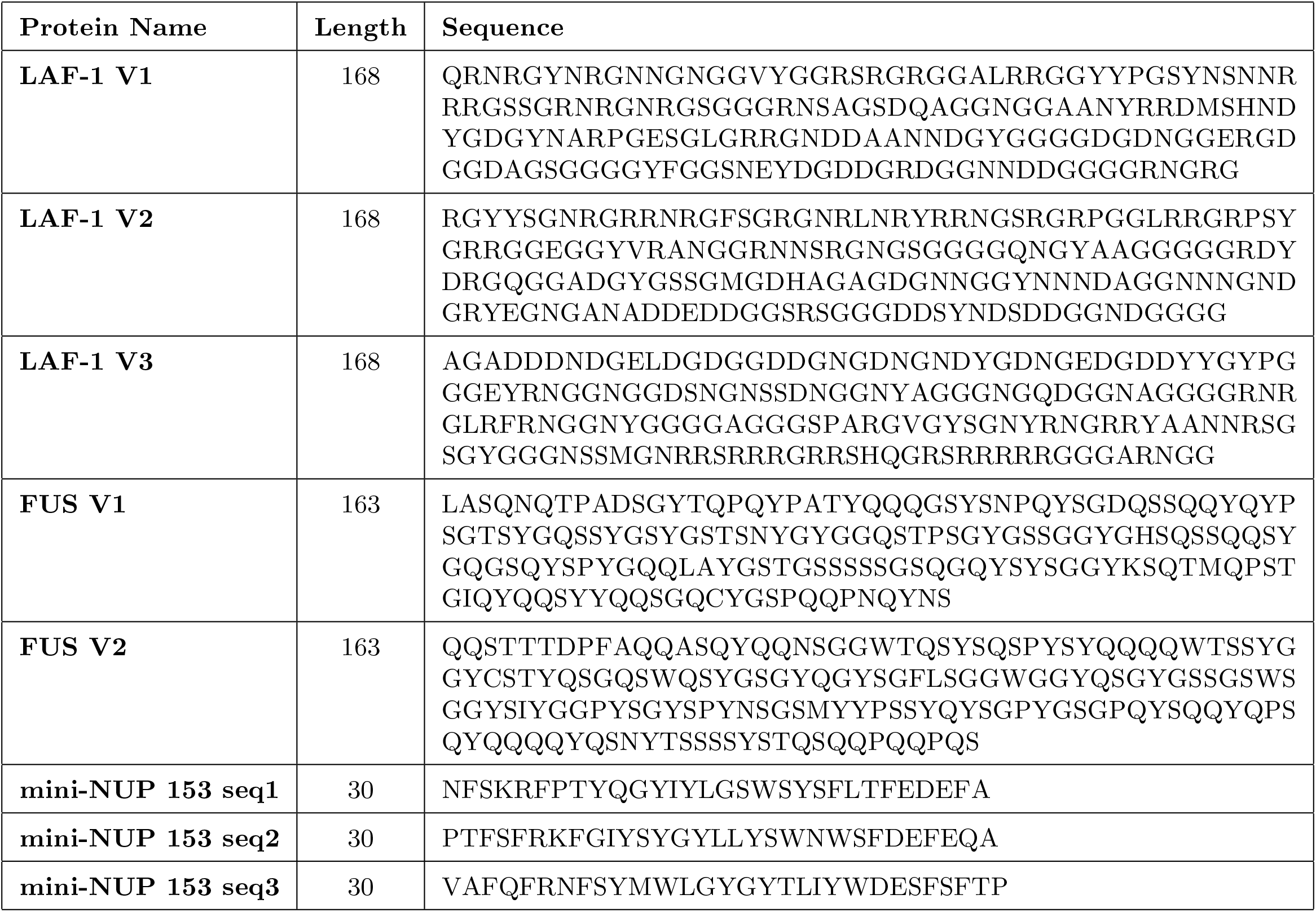
List of designed sequences used in this work in MD simulations of phase separation.

**Table 4:**
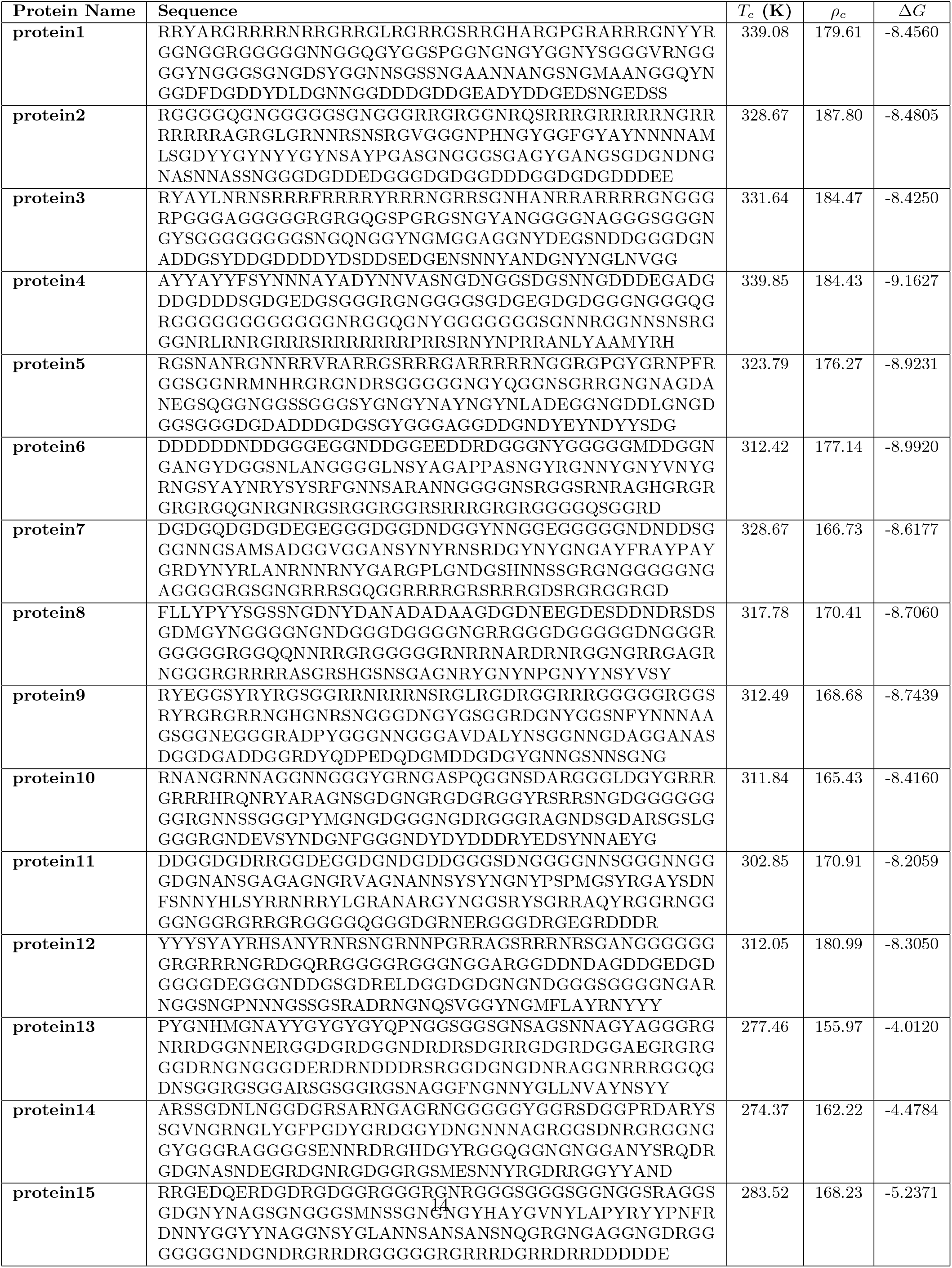
List of 15 additional designed LAF-1 RGG variants generated to span the (SCD, SHD) design space at fixed amino acid composition.

